# Preclinical Development of *Enterobacter ludwigii* LS57, a Human-Derived Swarming Strain, as a Live Biotherapeutic Product for Inflammatory Bowel Disease

**DOI:** 10.64898/2026.11.10.765554

**Authors:** Weijie Chen, Chaoyu Zhu, Yushan Guo, Weikang Yang, Yao Zhang, Zijie Qu, Mengya Wang, Shuai Gong, Shengdi Wu, Lingyu Zhu, Lihui Feng, Lei Liu

**Author notes:** **Correspondence: Lei Liu,****; Lihui Feng,**.

## Abstract

Inflammatory Bowel Disease (IBD) remains an area of major unmet need, and the undefined composition and manufacturing burden of fecal microbiota transplantation have motivated a shift toward single-strain Live Biotherapeutic Products (LBPs). Swarming bacteria are enriched in the stressed gut and limit mucosal damage, but no human-derived swarming strain has been developed as an LBP. We therefore report the complete preclinical development of *Enterobacter ludwigii* LS57, a swarming strain isolated from human feces. Chemistry, Manufacturing and Controls (CMC) covered a three-tier cell bank, 30 L fed-batch fermentation and a colon-targeted enteric-coated capsule, enabling consistent pilot-scale production. No treatment-related findings emerged in safety pharmacology (Sprague-Dawley rats) or in single-dose and Good Laboratory Practice (GLP)-compliant 1-week repeat-dose toxicity studies (Sprague-Dawley rats and Beagle dogs). LS57 was identified as *E. ludwigii* by 16S rRNA and whole-genome sequencing and produced metabolites with reported anti-inflammatory activity in culture. In Dextran Sulfate Sodium (DSS)-induced colitis, LS57 reduced weight loss, preserved colon length, lowered colonic pro-inflammatory cytokines and remodeled the gut microbiota. It also attenuated Trinitrobenzene Sulfonic Acid (TNBS)-induced colitis and enhanced the effect of ciclosporin A. *In vitro*, LS57 was non-hemolytic, non-cytotoxic, and acid-labile, formed no biofilm, and its resistance to aminopenicillins, cephamycins and imipenem was not transferable by conjugation. *In vivo*, LS57 colonized the inflamed colon without translocation in wild-type and interleukin-10-deficient mice, and experimental bacteremia was treatable with gentamicin or levofloxacin. LS57 combines efficacy across two mechanistically distinct colitis models with a safety package that addresses the principal regulatory concerns for LBPs, and a scalable manufacturing process, compatible with current good manufacturing practice, in which enteric coating overcomes the strain’s poor gastric survival. These data support clinical development of LS57 in IBD.

## Introduction

Inflammatory Bowel Disease (IBD), comprising Ulcerative Colitis (UC) and Crohn’s Disease (CD), is a chronic, relapsing inflammatory disorder of the gastrointestinal tract^1^. IBD has become a global health concern, with incidence rising 5-to 10-fold across Asia, Latin America, and Africa over the past three decades^2^, and the number of patients in Asia is projected to exceed 4 million by 2035^3^. Its pathogenesis reflects an interplay between genetic susceptibility, gut microbial dysbiosis, immune dysregulation, and environmental exposures such as dietary change and antibiotic overuse^4^. Symptoms including abdominal pain, diarrhea, and weight loss substantially impair quality of life, and the relapsing course frequently demands lifelong medication or surgery, imposing a heavy burden on patients and health systems^5^. Biologics and small-molecule drugs can induce and maintain remission but are not curative, and are limited by primary non-response, secondary loss of response, safety concerns, and cost^6^. A substantial proportion of patients therefore fail to achieve durable remission, underscoring the need for agents with distinct mechanisms^7^.

The gut microbiota’s central role in IBD pathogenesis has driven microbiota-targeted therapies^8^. Patients show reduced diversity, with depletion of *Bacteroidetes* and expansion of *Proteobacteria* such as *Escherichia coli*^9^, while loss of anti-inflammatory taxa such as *Faecalibacterium prausnitzii* correlates with disease severity^10^, and CD is further associated with adherent-invasive *E. coli* that sustains chronic immune stimulation^11^. Dysbiosis amplifies pro-inflammatory signalling through excessive TNF-α and IL-6 production and aberrant TLR and NOD2 sensing^12, 13^. A Live Biotherapeutic Product (LBP) is a biological product that contains live organisms, is intended for use in the prevention, treatment or cure of a human disease^14^. Fecal Microbiota Transplantation (FMT), the most clinically advanced such intervention, induces clinical and endoscopic remission in mild-to-moderate UC with a reassuring safety profile^15, 16^, but its wider adoption is constrained by donor selection, the absence of a defined active ingredient, batch-to-batch variability, and site-to-site standardization^17^. These limitations motivate a shift from undefined consortia to single-strain LBPs whose identity, potency and manufacture can be controlled.

Realizing that promise requires meeting four requirements that have no direct counterpart for conventional drugs, because the active substance replicates^14, 18^. The strain’s resistance profile and its potential to donate resistance genes to the resident microbiota must be defined^19, 20^. Translocation and bloodstream infection must be assessed, particularly in immunocompromised hosts^21, 22^. Manufacture must be scalable and reproducible, supported by a characterized cell bank^23, 24^. Toxicology must comply with Good Laboratory Practice (GLP) in a relevant species^19, 24^. Resistance transfer is the most difficult of these concerns to exclude: horizontal gene transfer can turn a commensal into a reservoir of resistance determinants^25, 26^, whereas breakpoint-based susceptibility testing characterises the candidate in isolation and says nothing about its capacity to donate resistance genes to other members of the gut community^26^. Candidate strains must therefore combine susceptibility testing^27^, genomic annotation of resistance determinants and mobile elements^28^, and conjugation assays where transfer is plausible^29^. Much of the LBP literature omits this package, hindering regulatory review and cross-study comparison^18, 19^.

Flagella-mediated motility has likewise been implicated in IBD pathogenesis: flagellin activates TLR5, drives NF-κB-dependent cytokine production and is a dominant antigen in CD^30, 31^. Swarming, a coordinated multicellular movement across surfaces, appears to act in the opposite direction^32, 33^. Reactive oxygen species, elevated luminal iron, and mucus disruption characteristic of the inflamed gut favor its emergence^33^, and using multiple species and their swarming-deficient mutants, we showed that only swarming-capable strains alleviate colitis, indicating that protection tracks the phenotype rather than species^32, 34^. Swarming bacteria lower intestinal oxygen tension, expand beneficial anaerobes and restore community homeostasis^32, 34^. That work, however, rested on *Enterobacter sp.* SM3, a mouse isolate. A murine strain cannot be developed as a human LBP, and whether a human-derived strain retains its phenotype and efficacy is unknown.

Here we report the isolation and complete preclinical development of *Enterobacter ludwigii* LS57, a swarming strain recovered from human feces. LS57 was selected for the swarming phenotype rather than for taxonomy, and its human origin addresses the central limitation of SM3. We show that LS57 retains swarming motility, attenuates colitis in two mechanistically distinct mouse models and enhances the effect of ciclosporin A. We then define its safety profile through *in vitro* characterization, *in vivo* translocation and antibiotic-treatability studies, and GLP-compliant toxicology in rats and dogs, and describe a scalable manufacturing process. These data constitute the preclinical package required to support an investigational new drug application for LS57.

## Results

### Isolation, Screening and Characterization of *E. ludwigii* LS57

To identify swarming-motile strains with therapeutic potential, we screened fecal samples from more than 100 human donors by swarming motility assay. LS57 was recovered as one of several swarming-positive isolates and was selected for development on the basis of phenotype rather than taxonomy. On Luria-Bertani (LB) agar at 37 °C for 18-24 h, LS57 formed white, raised colonies with a glistening, moist surface, smooth margins, and regular edges (Fig. 1A). Gram staining showed Gram-negative, short, plump bacilli with blunt ends, occurring singly or in pairs and without spores (Fig. 1B). In liquid LB at 37 °C, LS57 entered logarithmic growth within 1 h, remained in logarithmic phase to 4 h and reached stationary phase thereafter (Fig. 1C). On 0.5% agar at 37 °C and 40% relative humidity, colonies expanded radially and produced an irregular, diffuse film at the periphery characteristic of swarming; time-lapse imaging over 16 h showed the swarm reaching the edge of the plate (Fig. 1D).

**Figure 1.**
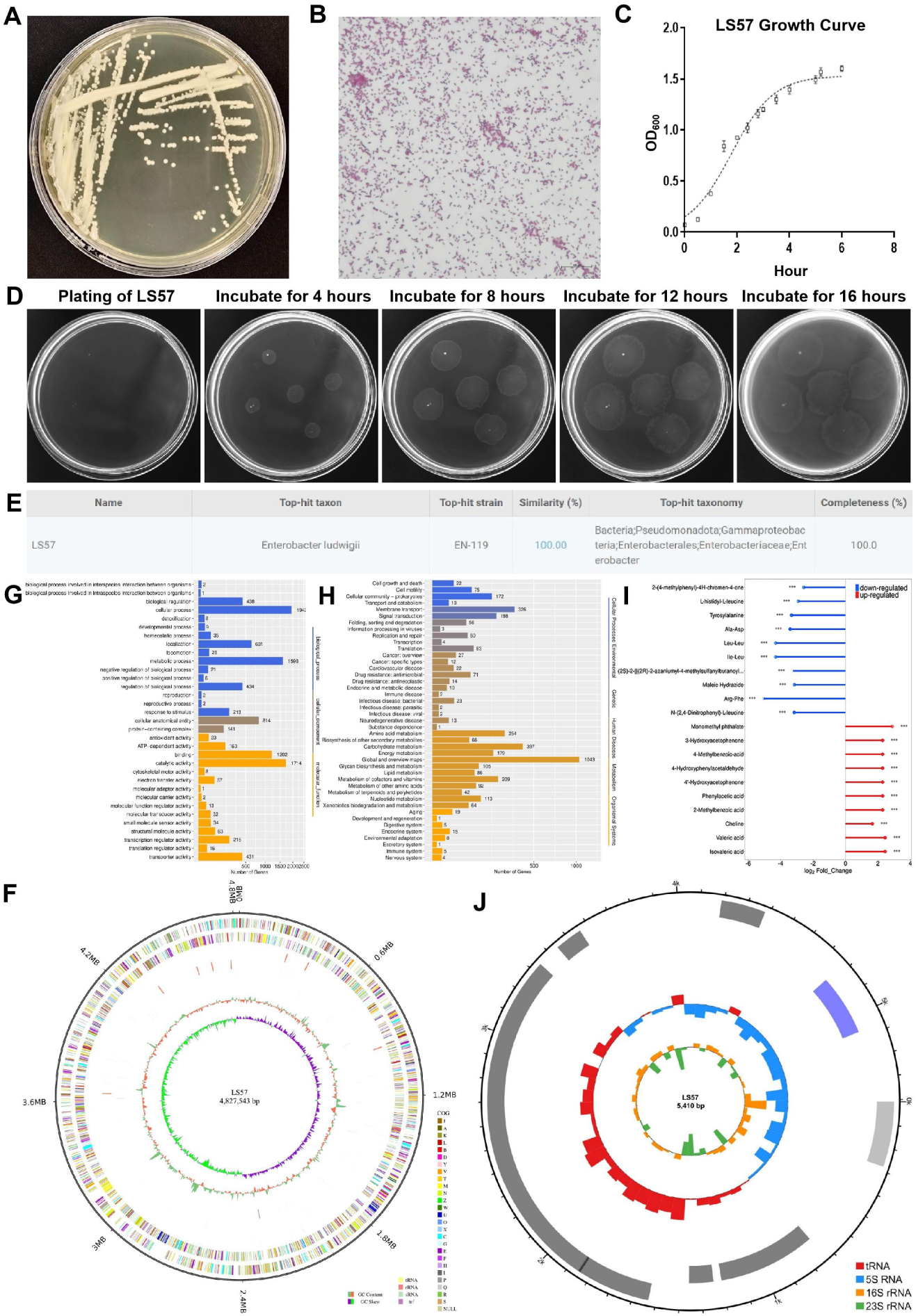
Isolation, screening and characterization of *E. ludwigii* LS57. (A) Colony morphology on LB agar. (B) Gram staining. (C) Growth curve in LB medium. (D) Swarming motility on 0.5% agar, photographed 0, 4, 8, 12, and 16 h after inoculation. (E) 16S rRNA gene-based identification of LS57. (F) Circular map of the 4,827,543 bp chromosome. (G) GO enrichment of LS57 genes. (H) KEGG pathway enrichment. (I) Anti-inflammatory metabolites in LS57 culture supernatant detected by LC-MS/MS. (J) Physical map of the 5,410 bp plasmid.

16S rRNA gene sequencing assigned LS57 to *Enterobacter ludwigii*, with 100% identity to the type-strain sequence in EzBioCloud^35^ (Fig. 1E). Because members of the *Enterobacter cloacae* complex are not resolved by 16S rRNA alone^36^, we completed the genome assembly and compared it with type strains of the complex. The assembly is a single circular chromosome of 4,827,543 bp with a G+C content of 54.6%, 4,571 predicted coding sequences of which 4,541 (99.34%) were assigned a function, and 153 non-coding RNAs accounting for 1.0% of the genome (Fig. 1F). Average Nucleotide Identity (ANI) and digital DNA–DNA Hybridization (dDDH), calculated with OrthoANIu^37^ and GGDC^38^, were highest against *E. ludwigii* EN-119 (GCF_001750725.1) at 98.94% and 91.9%, both above the species demarcation thresholds of 95%^39^ and 70%^40^ (Table 1). All other type strains of the complex, including *E. asburiae*, *E. bugandensis*, *E. cloacae*, *E. kobei* and *E. hormaechei*, fell to no more than 88.64% ANI and 35.9% dDDH.

**Table 1.** ANI and dDDH values of *E. ludwigii* LS57 and type strains of the *E. cloacae* complex.

| **Strain** | **Type strain of the *E. cloacae* complex** | **RefSeq** | **ANI (%)** | **dDDH (%)** |
|---|---|---|---|---|
| LS57 | *Enterobacter ludwigii* EN-119 | GCF_001750725.1 | 98.94 | 91.9 |
|  | *Enterobacter asburiae* 17Nkhm-UP2 | GCF_007035805.1 | 88.64 | 35.9 |
|  | *Enterobacter bugandensis* EB-247 | GCF_900324475.1 | 88.14 | 34.8 |
|  | *Enterobacter cloacae* 1382 | GCF_905331265.2 | 87.86 | 34.3 |
|  | *Enterobacter kobei* 11778-yvys | GCF_023023125.1 | 87.84 | 34.3 |
|  | *Enterobacter hormaechei* FDAARGOS 1433 | GCF_019048245.1 | 86.45 | 31.5 |

We next annotated the 4,571 predicted proteins against the Gene Ontology (GO)^41^ and KEGG^42^ databases to define the functional repertoire of the strain. GO terms were assigned to 3,186 genes (69.7% of the coding sequences), comprising 955, 3,974, and 5,368 assignments to cellular component, molecular function, and biological process, respectively (Fig. 1G), and KEGG assigned 3,394 genes (74.25%), among which metabolism was the dominant category (Fig. 1H). Liquid Chromatography-Tandem Mass Spectrometry (LC-MS/MS) of culture supernatants, relative to uninoculated medium, resolved 19 differentially abundant features, 10 enriched and 9 depleted in the spent culture (Fig. 1I). The depleted features were almost exclusively dipeptides (Leu-Leu, Ile-Leu, Arg-Phe, Ala-Asp, tyrosylalanine, and L-histidyl-L-leucine), consistent with peptide consumption during growth rather than a specific activity. Among the enriched features, 5 carry reported anti-inflammatory activity: choline, valeric acid, isovaleric acid, 4-hydroxyphenylacetaldehyde and 4′-hydroxyacetophenone (log2 fold change 1.5-2.4). In a related *E. ludwigii* isolate, choline-mediated immune tolerance protected mice against colitis^43^, while valeric and isovaleric acid belong to the Short-Chain Fatty Acid (SCFA) pool whose anti-inflammatory and barrier-protective actions in the colon are well documented^44^.

Assembly resolved a single circular plasmid of 5,410 bp in addition to the chromosome (Fig. 1J). The plasmid is small and carries 8 coding sequences, and no intact prophage was predicted in either replicon. The genome is thus closed and fully resolved, which allowed every subsequent safety endpoint to be interpreted against a complete sequence rather than a draft assembly.

### Therapeutic Efficacy of *E. ludwigii* LS57 in DSS-Induced Colitis

Dextran Sulfate Sodium (DSS) disrupts the intestinal barrier and activates innate immune responses, and it remains the most widely used means of inducing ulcerative-colitis-like injury in mice^45^. Having shown that murine swarming strains ameliorate DSS colitis^32^, we asked whether a human-derived swarming strain would do the same. Mice received 2.5% DSS in drinking water for 12 days and were gavaged daily with 8.0×10^8^ Colony-Forming Units (CFU) of LS57 (DSS+LS57) or an equal volume of LB medium (DSS+LB), with untreated mice given plain water as a healthy reference (No DSS+LB). Body weight was recorded daily and the experiment was terminated when body weight loss in the DSS+LB group exceeded 20%, followed by euthanasia. Healthy controls held their weight, whereas both DSS groups began to lose weight from day 6 and diverged progressively from day 7 onwards. By day 10 the DSS+LB mice had lost 15% of their starting weight against 10% in the DSS+LS57 group, and the gap widened to 16% versus 9% on day 11, both time points reaching significance (Fig. 2A). Colon length, measured at necropsy on day 12, was greater in LS57-treated mice than in DSS+LB controls (Fig. 2B). Hematoxylin and Eosin (H&E) staining showed extensive ulceration, crypt loss and architectural disruption in the DSS+LB colon, whereas LS57-treated tissue retained crypt architecture with damage confined to focal epithelial lesions (Fig. 2C).

**Figure 2.**
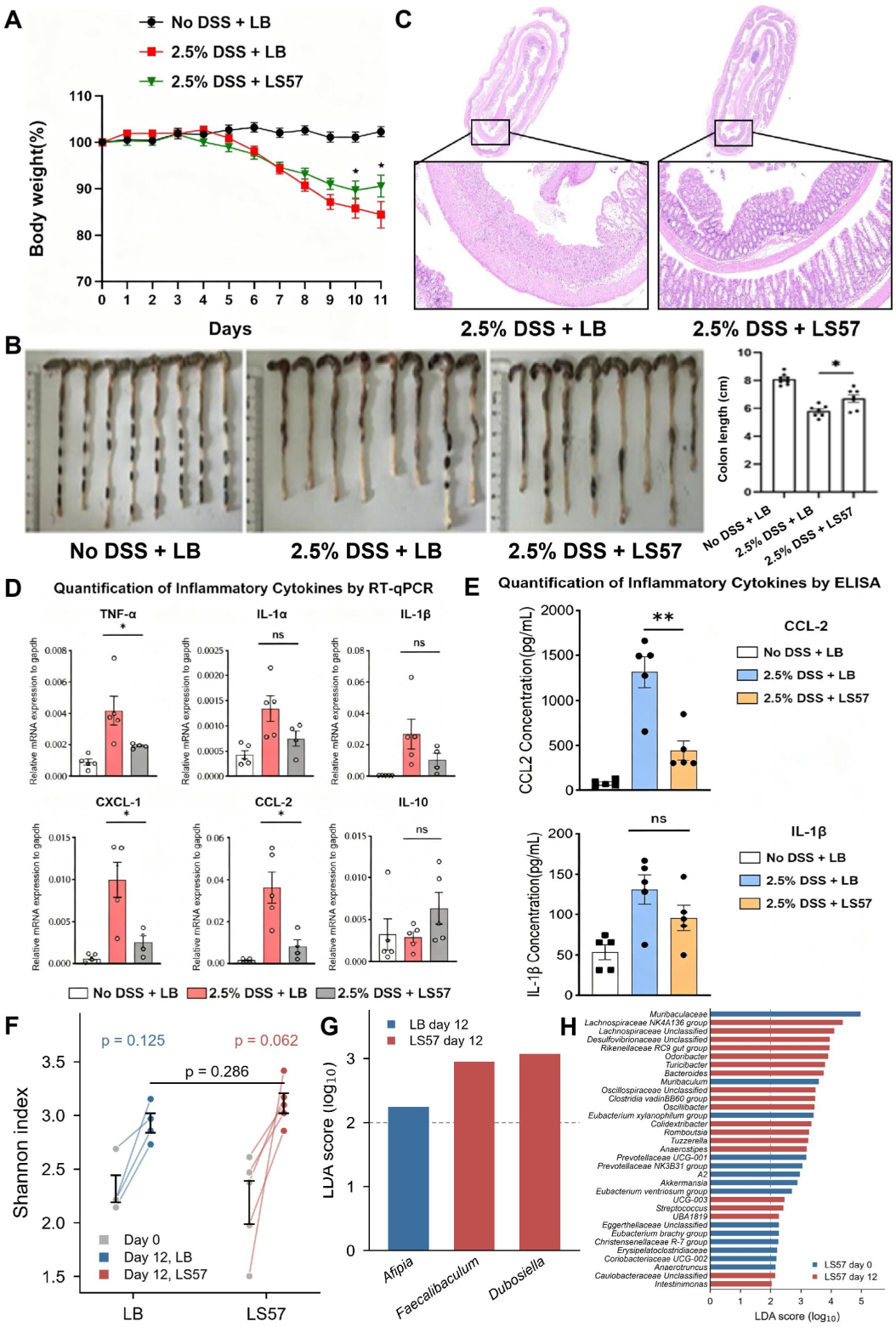
Therapeutic efficacy of *E. ludwigii* LS57 in DSS-induced colitis. (A) Body weight (% of day 0) of mice given plain water (No DSS+LB), 2.5% DSS with LB (DSS+LB) or 2.5% DSS with LS57 (DSS+LS57). (B) Representative colons and colon length measured at necropsy on day 12. (C) Representative H&E-stained colonic sections, with insets showing higher magnification of the boxed regions. (D) Relative mRNA expression of TNF-α, IL-1α, IL-1β, CXCL-1, CCL-2, and IL-10 in colon tissue, normalized to Gapdh; brackets compare DSS+LB with DSS+LS57. (E) CCL-2 and IL-1β concentrations in colon tissue (pg/mL). (F) Shannon index before (day 0) and after (day 12) DSS exposure. (G) LEfSe comparison of LB day 12 with LS57 day 12. (H) LEfSe comparison of LS57 day 0 with day 12.

To define the inflammatory signature behind this protection, colon tissue was analyzed at both transcript and protein level. Relative to healthy controls, DSS+LB mice showed higher mRNA for TNF-α, CXCL-1, and CCL-2 by RT-qPCR^46^, confirming induction of the mucosal inflammatory program (Fig. 2D). LS57 treatment lowered all three transcripts relative to DSS+LB, whereas IL-1α, IL-1β and IL-10 did not differ significantly between the two DSS groups, with IL-10 trending upward in LS57-treated tissue. At the protein level, ELISA^47^ showed that colonic CCL-2 was lower in LS57-treated mice than in DSS controls, whereas IL-1β decreased without reaching significance (Fig. 2E). LS57 therefore limited weight loss, colon shortening, architectural destruction, and cytokine induction within the same animals.

We next asked whether LS57 reshapes the gut microbiota during colitis. Fecal samples collected before DSS exposure (day 0) and on day 12 were profiled by 16S rRNA gene sequencing. The median Shannon index rose from 2.2 to 2.9 in LB controls and from 2.4 to 3.1 in LS57-treated mice, but the change was not significant within either group (Fig. 2F). The compositional differences described below are therefore not secondary to a loss of diversity. LEfSe^48^ separated the two groups at day 12: LB controls were characterized by *Afipia*, whereas the LS57 group was enriched for *Faecalibaculum* and *Dubosiella* (Fig. 2G). Within the LS57 group, day 12 relative to day 0 was marked by 19 enriched and 14 depleted genera (Fig. 2H). The enriched set included *Odoribacter*, *Rikenellaceae* RC9 gut group, *Turicibacter*, *Bacteroides*, *Colidextribacter*, *Romboutsia*, *Anaerostipes*, and *Lachnospiraceae* NK4A136 group, several of which are associated with SCFA production and mucosal homeostasis^44^. The depleted set included *Muribaculaceae*, *Muribaculum*, *Akkermansia*, *Prevotellaceae* UCG-001, and *Eubacterium xylanophilum* group.

### Therapeutic Efficacy of *E. ludwigii* LS57 in TNBS-Induced Colitis, Combination with Ciclosporin A, and *In Vivo* Biodistribution

Trinitrobenzene Sulfonic Acid (TNBS) dissolved in ethanol elicits a transmural, T-cell-driven colitis that reproduces the discontinuous inflammation and Th1 polarization of CD, and it therefore interrogates a mechanism distinct from the epithelial injury produced by DSS^49^. Mice were fasted for 16 h then received 2% TNBS in 40% ethanol rectally and were gavaged daily for 4 days with 7.0×10^8^ CFU of LS57 (TNBS+LS57) or an equal volume of LB medium (TNBS+LB). Both groups lost 16-18% of body weight within 24 h of instillation and recovered in parallel thereafter, with no significant difference at any time point (Fig. 3A). Colon length, measured once TNBS+LB mice had regained 95% of baseline weight, was greater in TNBS+LS57 mice (Fig. 3B). LS57 therefore attenuated TNBS-induced damage without modifying the systemic weight trajectory, the behavior expected of a strain that acts within the gut lumen.

**Figure 3.**
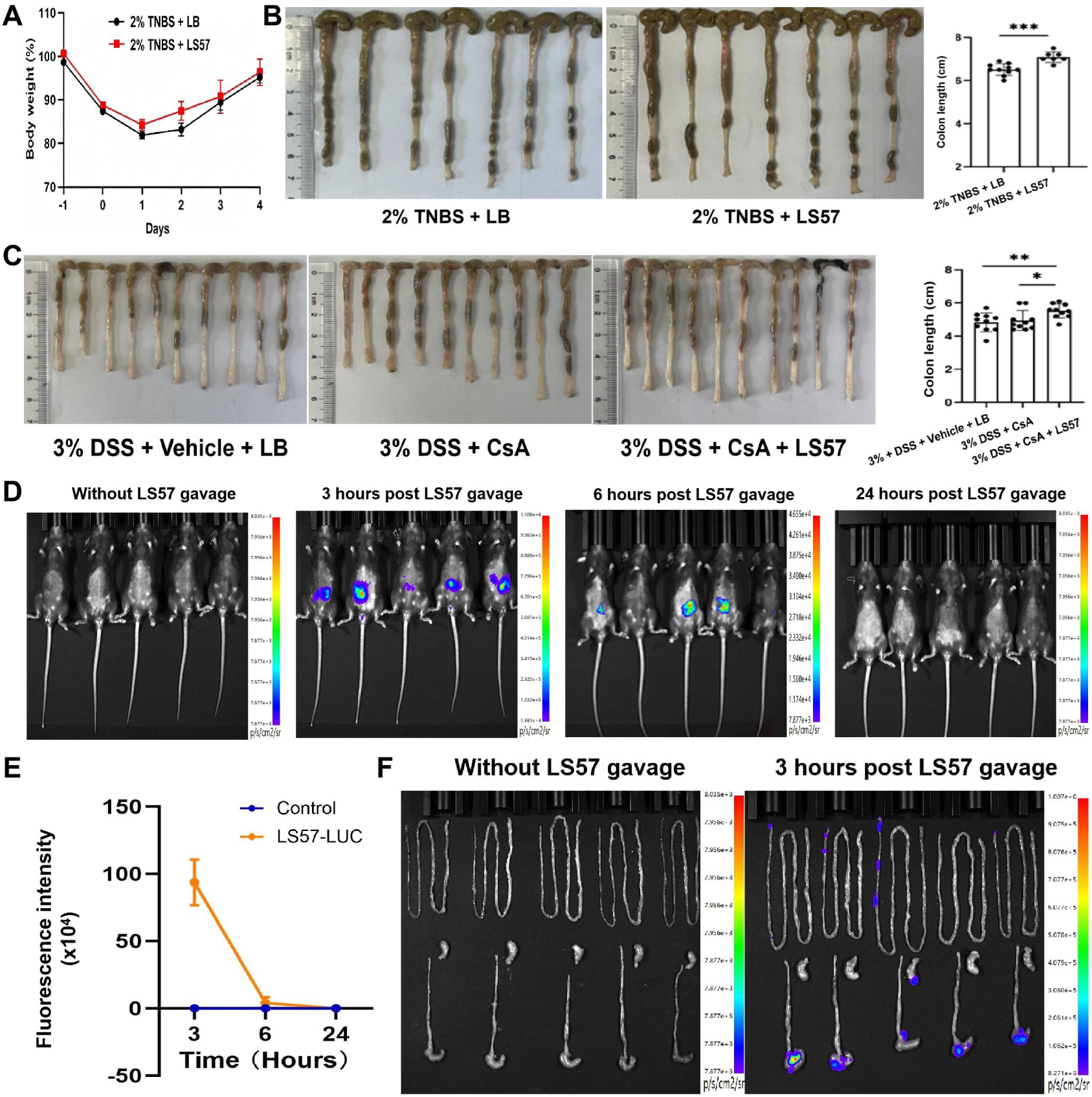
Therapeutic efficacy of *E. ludwigii* LS57 in TNBS-induced colitis, combination with ciclosporin A, and *in vivo* biodistribution. (A) Body weight (% of day 0) of mice given 2% TNBS in 40% ethanol and gavaged daily with LS57 (TNBS+LS57) or an equal volume of LB (TNBS+LB). (B) Representative colons and colon length at the end of the experiment. (C) Representative colons and colon length in mice given 3% DSS with vehicle plus LB (DSS+Vehicle+LB), CsA alone (DSS+CsA) or CsA plus LS57 (DSS+CsA+LS57). (D) Whole-body bioluminescence images before gavage and 3, 6 and 24 h after a single dose of LS57-LUC; the color scale is radiance in p/s/cm²/sr. (E) Abdominal bioluminescence quantified at the indicated times in mice given LS57-LUC or sterile LB. (F) *Ex vivo* bioluminescence of the explanted small intestine, cecum, and colon without gavage and 3 h after LS57-LUC.

Because a live biotherapeutic will be given alongside conventional immunosuppression rather than instead of it, we next asked whether LS57 augments ciclosporin A (CsA), the calcineurin inhibitor used for steroid-refractory IBD^50^. Colitis was induced with 3% DSS over 11 days, and mice received CsA alone (15 mg/kg/day; DSS+CsA), CsA plus LS57 (8.0×10^8^ CFU/day; DSS+CsA+LS57), or vehicle plus LB (DSS+Vehicle+LB). Colon length was comparable between the DSS+Vehicle+LB and DSS+CsA groups, but was significantly greater in DSS+CsA+LS57 than in either group (Fig. 3C). The combination therefore outperformed CsA alone rather than merely reproducing its effect, which is the behavior required of an adjunct intended to improve rather than replace immunosuppressive therapy.

To determine the *in vivo* biodistribution of LS57 following oral administration, the luxCDABE operon was integrated into the LS57 genome by Tn7 transposition to generate a bioluminescent reporter strain (LS57-LUC). Mice received a single 200 µL dose of LS57-LUC (4.0×10^9^ CFU/mL; 8.0×10^8^ CFU in total, the dose used in the colitis studies) or sterile LB by gavage, and bioluminescence and its distribution within the digestive tract were assessed before gavage and at 3, 6, and 24 h. Abdominal signal was detectable 3 and 6 h after gavage, declined progressively over the observation period, and was no longer detectable above background at 24 h (Fig. 3D-E). Upon removal of the gastrointestinal tissues, the remaining signal was found mainly in the small intestine and cecum 3 h after gavage (Fig. 3F).

### *In Vitro* Safety Assessment of *E. ludwigii* LS57

The stomach is the first barrier an orally administered strain must cross. LS57 was therefore resuspended in Simulated Gastric Fluid (SGF) at pH 2, 3, and 4, spanning the fasting-to-fed range of gastric pH, with cells held in 0.9% NaCl as the control. No LS57 was recovered at pH 2, and only a small fraction survived at pH 3 and pH 4 (Fig. 4A). LS57 is thus acid-labile, a limitation for delivery rather than a safety liability, since it restricts the proportion of an oral dose that survives gastric transit, consistent with the transient, gut-confined signal seen by imaging (Fig. 3D-F). An enteric-coated capsule, which protects the strain during gastric transit and releases it in the intestine, is therefore the preferred dosage form for clinical use.

**Figure 4.**
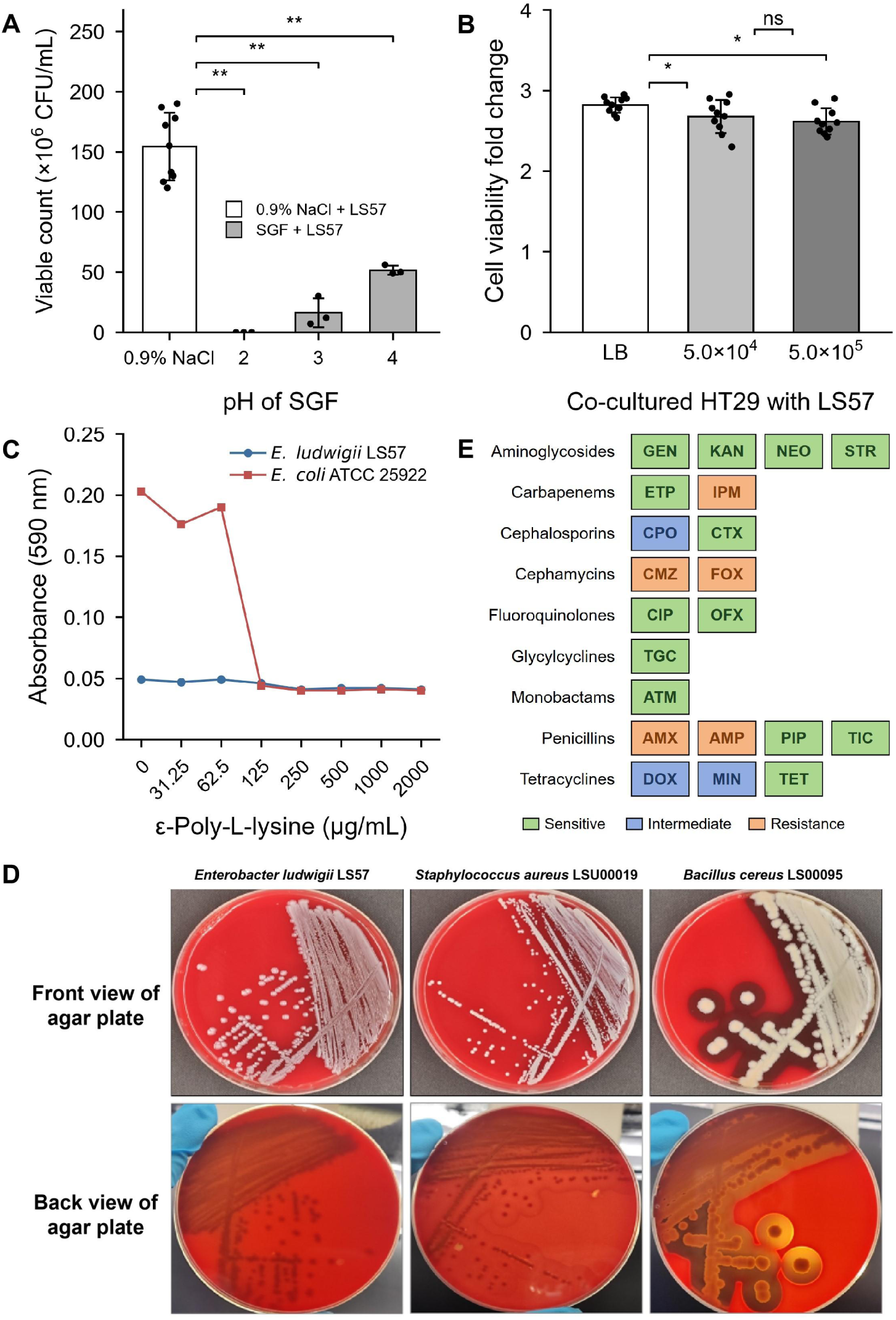
*In Vitro* Safety Assessment of *E. ludwigii* LS57. (A) Viability of LS57 (CFU/mL) after incubation in 0.9% NaCl (0.9% NaCl + LS57) or simulated gastric fluid (SGF + LS57) at pH 2, 3, and 4. (B) Viability of HT-29 cells after 24 h of co-culture with LS57 at 5.0×10^4^ or 5.0×10^5^ CFU/mL, or with an equal volume of LB, measured by CCK-8 and expressed as fold change relative to untreated cells. (C) Biofilm biomass (A_590_) of LS57 and *E. coli* ATCC 25922 after 24 h with ε-poly-L-lysine at the indicated concentrations. (D) Hemolysis on 5% defibrinated sheep blood agar after 24 h. (E) Antibiotic susceptibility of LS57 by disk diffusion for 21 agents across 9 classes; each cell gives the drug abbreviation and color gives the interpreted category.

Cytotoxicity was assessed in human colon cancer cells (HT-29) co-cultured for 24 h with LS57 at 5.0×10^4^ and 5.0×10^5^ CFU/mL, with an equal volume of LB as the control. Viability measured by CCK-8, expressed as fold change against untreated cells, was modestly but consistently lower in both LS57-treated groups than in the LB control, and the difference did not increase when the dose was raised tenfold (Fig. 4B). This small, non-dose-dependent difference is not consistent with a cytotoxic effect, and LS57 therefore shows no appreciable cytotoxicity towards intestinal epithelial cells over the dose range tested.

Biofilm formation was quantified by crystal violet staining after 24 h of growth in the presence of ε-poly-L-lysine, a cationic antimicrobial peptide used here as a biofilm inhibitor, across 0-2000 µg/mL. LS57 showed a consistently low A_590_ of 0.047±0.003 at every concentration tested, approximately fourfold lower than the 0.20 reached by *E. coli* ATCC 25922 in drug-free medium, and its adherence was unaffected across the entire range; in contrast, the control strain declined to the same level (A_590_ around 0.04) at concentrations of 125 µg/mL and above (Fig. 4C). LS57 therefore classifies as a non-biofilm-forming strain^51^, which argues against a biofilm-based persistence phenotype in the gut.

Hemolytic activity was assessed on 5% defibrinated sheep blood agar after 24 h, with *Staphylococcus aureus* LSU00019 (positive for α-hemolysis) and *Bacillus cereus* LS00095 (positive for β-hemolysis) as reference strains, and the plates were read from the front and the back. Both references cleared the agar, partially and completely respectively, whereas no zone formed around LS57 in either view, which is γ-hemolysis (Fig. 4D). The strain therefore lacks the membrane-damaging activity that defines the hemolysis virulence trait.

Genome annotation against the Comprehensive Antibiotic Resistance Database (CARD)^28^ identified 180 resistance genes distributed across 7 resistance mechanisms, dominated by antibiotic efflux (101 genes) and target alteration (64 genes), and predicted resistance to 33 drug classes (Supplementary Table S1). Most of these determinants are intrinsic chromosomal features of the *E. cloacae* complex, such as efflux systems, a chromosomal ampC, and porin-related permeability barriers, rather than acquired mobile elements. To test these predictions phenotypically, 21 agents across 9 classes were examined by disk diffusion^52^ (Table 2) and a further 7 agents across 6 classes by Minimum Inhibitory Concentration (MIC)^53^ determination (Table 3), with the cognate resistance genes identified for each agent listed alongside. Disk diffusion classified 5 agents as Resistant (R), 3 as Intermediate (I), and 13 as Sensitive (S) (Fig. 4E, Supplementary Fig. S1A). In the MIC panel, values ranged from 7.81 µg/mL for iclaprim to >1000 µg/mL for pleuromutilin (Table 3). These agents were selected to interrogate specific determinants predicted by genome annotation, rather than for clinical susceptibility interpretation. The combined pattern (cefotaxime active, whereas cephamycins, imipenem and the aminopenicillins were inactive) is characteristic of chromosomal ampC expression together with reduced outer-membrane permeability^54^, both intrinsic to the *E. cloacae* complex, rather than of an acquired plasmid-borne determinant. The same profile explains why gentamicin and levofloxacin, the two agents that controlled bacteremia *in vivo*, were among those to which LS57 was susceptible.

**Table 2.** Antibiotic susceptibility results of *E. ludwigii* LS57 determined by disk diffusion.

| **Antibiotic** |  | **Inhibition zone diameter (mm)** | **Interpretation standard (mm)** |  |  | **Interpretation** |
|---|---|---|---|---|---|---|
| **Class** | **Agent** |  | **Resistant (R)** | **Intermediate (I)** | **Sensitive (S)** |  |
| Aminoglycosides | Gentamicin | 21.0±0.5 | ≤12.0 | 13.0-14.0 | ≥15.0 | S |
|  | Kanamycin | 19.5±0.5 | ≤13.0 | 14.0-17.0 | ≥18.0 | S |
|  | Neomycin | 20.0±1.0 | ≤12.0 | 13.0-16.0 | ≥17.0 | S |
|  | Streptomycin | 18.0±0.5 | ≤11.0 | 12.0-14.0 | ≥15.0 | S |
| Carbapenems | Ertapenem | 22.0±2.0 | ≤18.0 | 19.0-21.0 | ≥22.0 | S |
|  | Imipenem | 19.0±3.0 | ≤19.0 | 20.0-22.0 | ≥23.0 | R |
| Cephalosporins | Cefpirome | 20.0±1.0 | ≤19.0 | 20.0-22.0 | ≥23.0 | I |
|  | Cefotaxime | 27.0±2.0 | ≤22.0 | 23.0-25.0 | ≥26.0 | S |
| Cephamycins | Cefmetazole | 0.0±0.0 | ≤12.0 | 13.0-15.0 | ≥16.0 | R |
|  | Cefoxitin | 0.0±0.0 | ≤14.0 | 15.0-17.0 | ≥18.0 | R |
| Fluoroquinolones | Ciprofloxacin | 28.5±0.5 | ≤15.0 | 16.0-20.0 | ≥21.0 | S |
|  | Ofloxacin | 22.0±0.5 | ≤12.0 | 13.0-15.0 | ≥16.0 | S |
| Glycylcyclines | Tigecycline | 19.5±0.5 | ≤12.0 | 13.0-15.0 | ≥16.0 | S |
| Monobactams | Aztreonam | 32.0±3.0 | ≤17.0 | 18.0-20.0 | ≥21.0 | S |
| Penicillins | Amoxicillin | 0.0±0.0 | ≤13.0 | 14.0-17.0 | ≥18.0 | R |
|  | Ampicillin | 0.0±0.0 | ≤13.0 | 14.0-16.0 | ≥17.0 | R |
|  | Piperacillin | 22.0±2.0 | ≤17.0 | 18.0-20.0 | ≥21.0 | S |
|  | Ticarcillin | 21.0±1.0 | ≤14.0 | 15.0-19.0 | ≥20.0 | S |
| Tetracyclines | Doxycycline | 13.5±0.5 | ≤12.0 | 13.0-15.0 | ≥16.0 | I |
|  | Minocycline | 16.0±0.5 | ≤14.0 | 15.0-18.0 | ≥19.0 | I |
|  | Tetracycline | 18.0±0.2 | ≤11.0 | 12.0-14.0 | ≥15.0 | S |

**Table 3.**
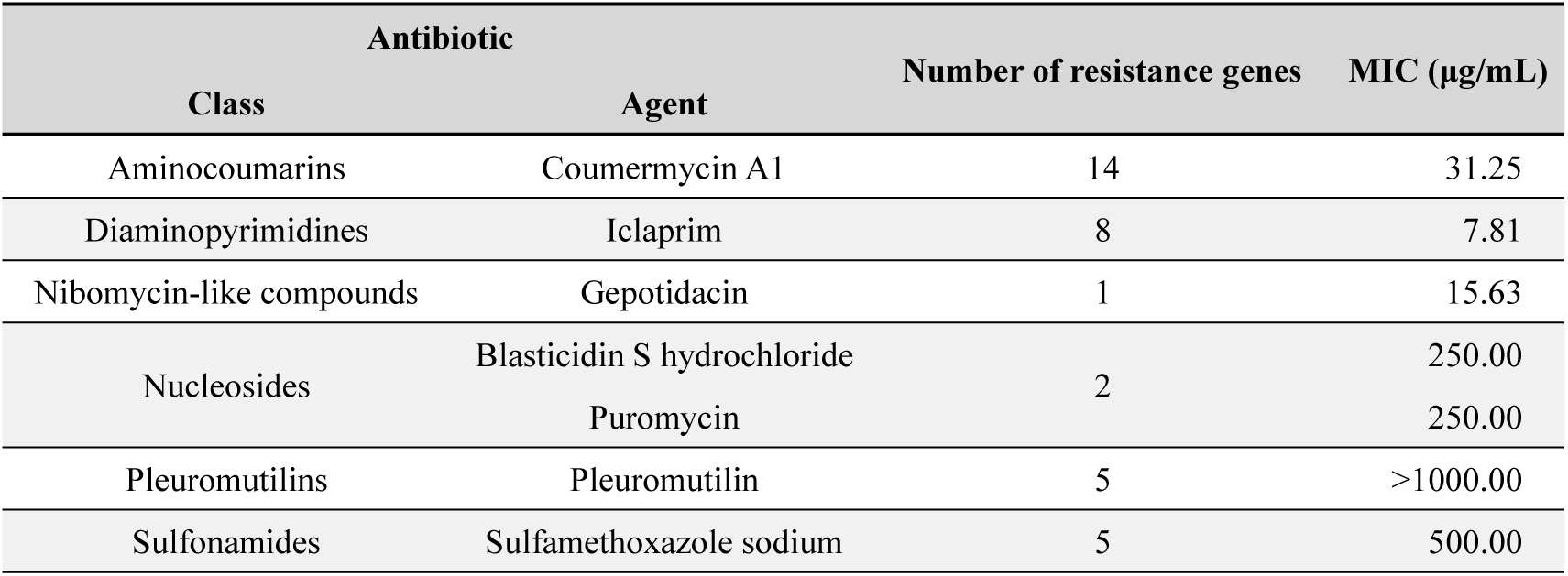
Antibiotic susceptibility results of *E. ludwigii* LS57 determined by MIC assay.

| **Antibiotic** |  | **Number of resistance genes** | **MIC (μg/mL)** |
|---|---|---|---|
| **Class** | **Agent** |  |  |
| Aminocoumarins | Coumermycin A1 | 14 | 31.25 |
| Diaminopyrimidines | Iclaprim | 8 | 7.81 |
| Nibomycin-like compounds | Gepotidacin | 1 | 15.63 |
| Nucleosides | Blasticidin S hydrochloride | 2 | 250.00 |
|  | Puromycin |  | 250.00 |
| Pleuromutilins | Pleuromutilin | 5 | >1000.00 |
| Sulfonamides | Sulfamethoxazole sodium | 5 | 500.00 |

The resistance genotype therefore exceeds the phenotype, and the safety-relevant question is not the resistance profile of LS57 itself but whether any of these determinants can be mobilised. Mobility was assessed both bioinformatically and experimentally. LS57 harbours a single 5,410 bp plasmid with 8 coding sequences, none of which is an Antibiotic Resistance Gene (ARG), so ARGs cannot be disseminated from this strain by plasmid transfer. Six Insertion Sequences (ISs) were annotated within or adjacent to ARG-containing regions; all six are truncated, sharing only 25.0-48.8% nucleotide identity with their closest reference elements, and no full-length copy was present in the genome, arguing against the formation of functional composite transposons (Supplementary Table S2). One region flagged as prophage-like by Prophage Hunter^55^ was not supported as a phage by PHASTEST^56^, consistent with a defective remnant. Mobilisation was then tested directly in an *in vitro* conjugation assay with *E. coli* MG1655 (pUC57-Tac-mCherry-Gen) as recipient; an RP4-carrying Stbl3 donor served as positive control and transferred at 65.1% (175/269 recipient cells), whereas no transconjugant was recovered from LS57 under selection for doxycycline or cefmetazole resistance (0/269; Supplementary Fig. S1B), placing its transfer frequency below the detection limit of the assay. Together, the genetic and functional evidence indicates that the resistance determinants of LS57 are not readily mobilisable and that the risk of horizontal transfer to the resident gut microbiota is low.

### *In Vivo* Safety Assessment of *E. ludwigii* LS57

The capacity of LS57 to disseminate beyond the gut was assessed in DSS colitis, the condition in which barrier disruption makes systemic spread most likely. C57BL/6 Wild-Type (WT) and Interleukin-10-KnockOut (IL-10 KO) mice^21^ received 2.5% DSS for 10 days and were gavaged daily with 8.0×10^8^ CFU of LS57 or *E. coli* Nissle 1917 (EcN)^22^; at necropsy, colon, Mesenteric Lymph Nodes (MLNs), blood, liver and spleen were cultured, with species identity confirmed by Polymerase Chain Reaction (PCR). In WT mice, both strains colonized the inflamed colon (LS57 80.0%, EcN 100.0%) and were recovered from MLNs at comparable frequencies (42.9% and 57.1%), but neither was recovered from blood, liver or spleen in any animal (Fig. 5A). IL-10 KO mice showed the same pattern of containment: LS57 colonized more consistently than EcN (83.3% vs 16.7%), with correspondingly higher MLN recovery (28.6% vs 14.3%), and again no dissemination to blood, liver or spleen (Fig. 5B, Supplementary Fig. S2). Assay sensitivity was confirmed by blood-culture spike-in controls that consistently yielded 180-200 colonies per plate. In contrast, untreated DSS-only IL-10 KO mice developed sporadic bacteremia with endogenous organisms (*Shigella flexneri*, *Escherichia fergusonii*, and *Enterococcus faecalis*, identified by 16S rRNA sequencing), whereas neither treatment group did. Colonization with LS57 was therefore not associated with increased translocation, whereas untreated colitis was, suggesting that LS57, like EcN, protects against translocation of the resident microbiota.

**Figure 5.**
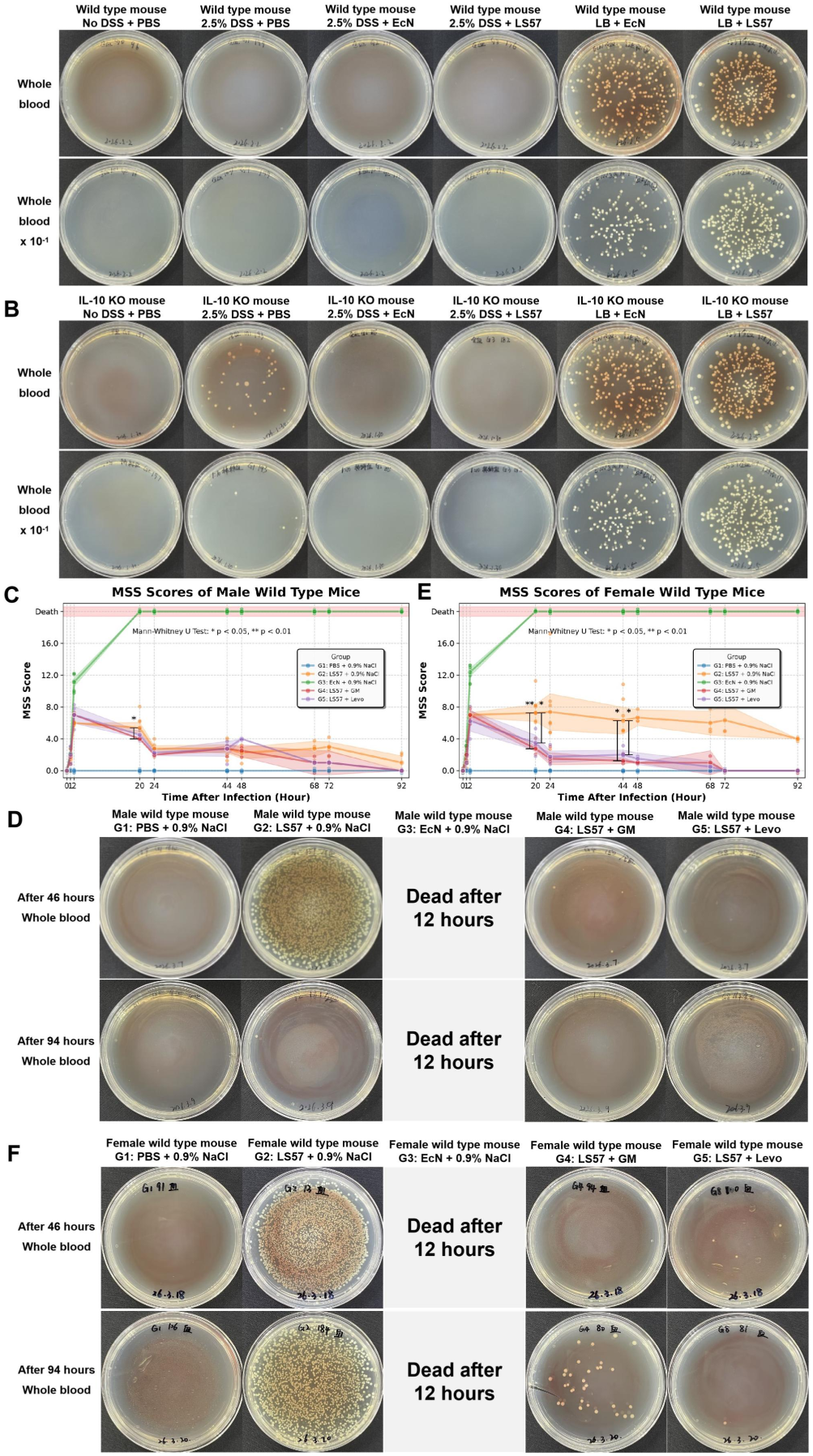
In Vivo Safety Assessment of *E. ludwigii* LS57. (A) Blood cultures from DSS-treated WT mice given daily LS57 or EcN, with DSS-only and untreated controls; all were negative for both strains. (B) Blood cultures from DSS-treated IL-10 KO mice given daily LS57 or EcN; cultures were negative for both strains, whereas DSS-only IL-10 KO controls yielded sporadic endogenous bacteremia. (C) MSS score of male mice after intraperitoneal infection with 1.0×10^8^ CFU LS57 and treatment with GM or Levo. (D) Blood cultures of the same male mice at 46 and 94 h after infection. (E) MSS score of female mice after intraperitoneal infection with 1.0×108 CFU LS57 and treatment with GM or Levo. (F) Blood cultures of the same female mice at 46 and 94 h after infection.

Whether LS57 infection can be controlled by standard antibiotics was examined in a murine bacteremia model. WT mice were inoculated intraperitoneally with 1.0×10^8^ CFU of LS57 and, once sepsis was confirmed by a rising Murine Sepsis Score (MSS)^57^ and positive blood cultures, were treated from 2 h post-infection with gentamicin (GM, 62 mg/kg every 24 h, intramuscular) or levofloxacin (Levo, 41 mg/kg every 12 h, intravenous), at doses scaled from human clinical regimens. Blood, liver, spleen, kidney and peritoneal lavage fluid were sampled at 46 and 94 h post-infection, and MSS was scored at 0, 1, 2, 20, 24, 44, 48, 68, 72, and 92 h. In males, GM reduced MSS relative to untreated controls at 20 h, whereas Levo did not differ from controls at any time point (Fig. 5C); blood cultures were sterile under both regimens at 46 h and remained sterile at 94 h (Fig. 5D). In females, both agents reduced MSS at 20 h and again at 44 h (Fig. 5E); blood cultures were sterile at 46 h, but low numbers of colonies reappeared at 94 h (Fig. 5F). Tissue clearance lagged behind clearance from blood: at 46 h, colonies were still recovered from liver, spleen, and kidney in at least one treated group; by 94 h these organs were sterile in males, whereas females retained low colony numbers in liver and spleen, and peritoneal lavage fluid remained positive in males (Supplementary Fig. S3) and females (Supplementary Fig. S4). An equal inoculum of EcN, tested in parallel, caused 100% mortality within 12 h. Thus, even when introduced at a high inoculum by the intraperitoneal route, LS57 caused no mortality in treated or untreated animals and was cleared from blood by both standard regimens, although sterilisation of deep tissues was slower and differed between sexes.

### CMC and GLP Toxicity Evaluation of *E. ludwigii* LS57 Drug Substance and Drug Product

To support industrial development and regulatory registration, a full Chemistry, Manufacturing and Controls (CMC) program, conducted in line with pharmacopeial standards and current Good Manufacturing Practice (cGMP) principles, was completed for the single-strain *E. ludwigii* LS57, beginning with the standardized construction of a three-tier cell bank system consisting of the Primary, Master, and Working Cell Banks (PCB, MCB, and WCB). Each tier was characterized by 16S rRNA species homology analysis, viable count calibration, strain purity testing, screening for exogenous microbes and pathogenic bacteria, and *in vivo* toxicity assays. Passage control met regulatory requirements for live microecological products: no more than 4 generations elapse between PCB initiation and fermentation harvest, and no more than 2 generations between WCB recovery and harvest. Genetic stability was further confirmed over 30 consecutive subcultures, with no detectable alteration in morphological, biochemical or growth phenotypes. All results met the predetermined internal quality standards, providing a traceable and genetically stable seed foundation for process development and large-scale production.

Based on the qualified production seed system, a complete bulk manufacturing process, comprising seed expansion, 30 L fed-batch fermentation, bacterial harvesting and washing, stabilizer resuspension, vacuum freeze-drying, and powder preparation, was established through fermentation medium screening, iterative optimization of critical process parameters and laboratory-scale verification. The finalized gradient fed-batch strategy initiated feed supplementation once the broth reached an OD_600_ of at least 18.0, with harvest at an OD_600_ of 45.0-55.0; three consecutive 30 L pilot batches showed consistent yield, viability and key quality attributes. To protect viable bacteria against gastric inactivation during oral delivery, excipient compatibility screening, and formulation optimization across seven prototype formulations yielded a pH-dependent colon-targeted enteric capsule, in which lyophilized LS57 powder blended with corn starch and lactose (7:3, w/w) is filled into enteric gelatin capsules that remain intact for 2 h in simulated gastric fluid (pH 1.2) and 3 h in simulated small intestinal fluid (pH 6.8) but disintegrate within 1 h in simulated colonic fluid (pH 7.8). Quality specifications (Table 4), with all core analytical methods validated for specificity, precision, and accuracy, were established for both the Drug Substance (DS, lyophilized bacterial powder) and the Drug Product (DP, enteric-coated capsule formulation), and the stability of both was investigated under long-term, accelerated and stress conditions; after 12 months at −20 °C, all critical quality attributes of the three pilot batches remained within specification, supporting a preliminary shelf life of 12 months under frozen storage.

**Table 4.**
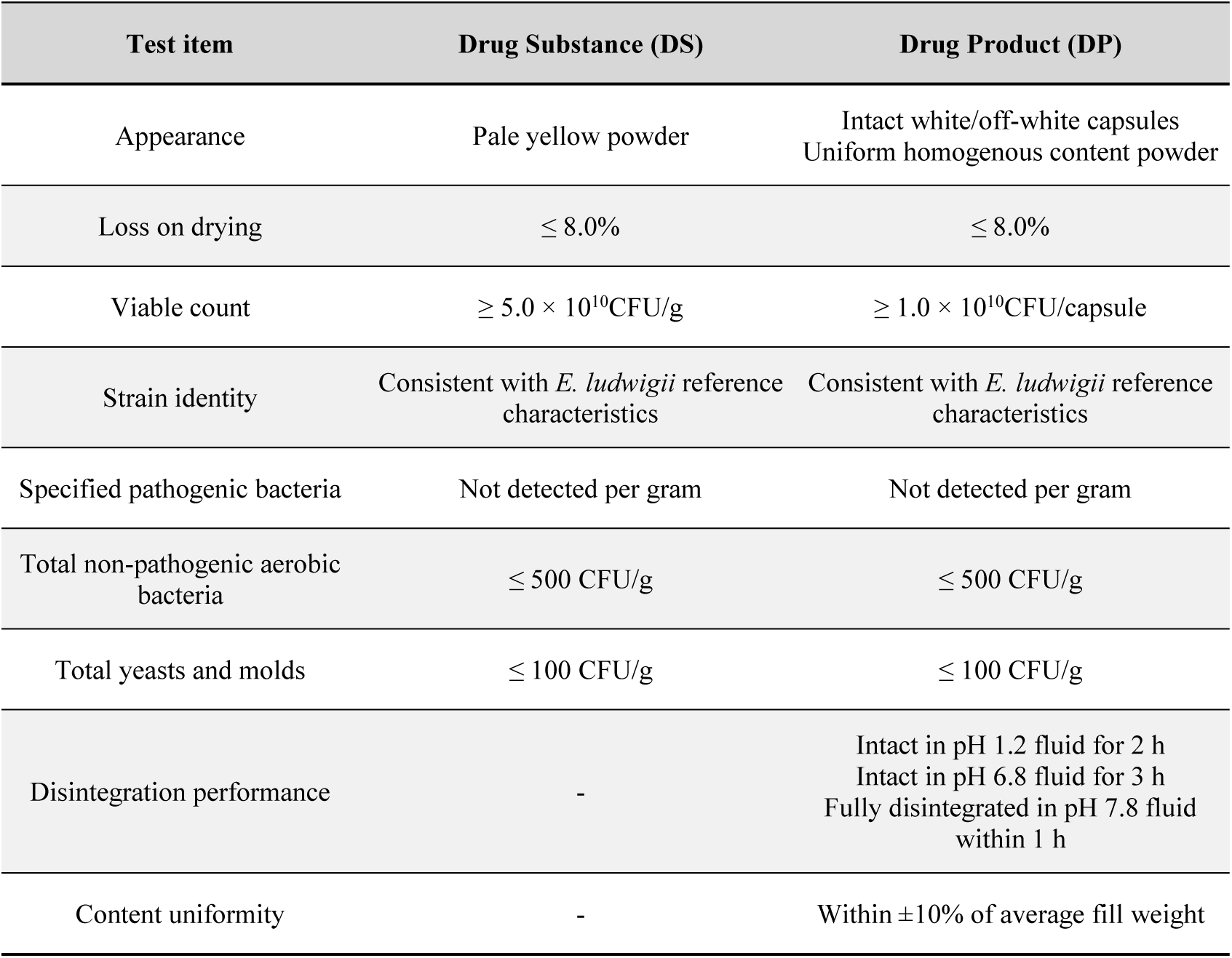
Quality specifications for *E. ludwigii* LS57 drug substance and drug product.

| **Test item** | **Drug Substance (DS)** | **Drug Product (DP)** |
|---|---|---|
| Appearance | Pale yellow powder | Intact white/off-white capsules<br>Uniform homogenous content powder |
| Loss on drying | ≤ 8.0% | ≤ 8.0% |
| Viable count | ≥ 5.0 × 10<sup>10</sup>CFU/g | ≥ 1.0 × 10<sup>10</sup>CFU/capsule |
| Strain identity | Consistent with *E. ludwigii* reference characteristics | Consistent with *E. ludwigii* reference characteristics |
| Specified pathogenic bacteria | Not detected per gram | Not detected per gram |
| Total non-pathogenic aerobic bacteria | ≤ 500 CFU/g | ≤ 500 CFU/g |
| Total yeasts and molds | ≤ 100 CFU/g | ≤ 100 CFU/g |
| Disintegration performance | - | Intact in pH 1.2 fluid for 2 h<br>Intact in pH 6.8 fluid for 3 h<br>Fully disintegrated in pH 7.8 fluid within 1 h |
| Content uniformity | - | Within ±10% of average fill weight |

Safety pharmacology was evaluated in Sprague-Dawley (SD) rats following a single oral gavage administration of LS57 at dose levels of 1.2×10^10^, 1.2×10^11^, and 6.0×10^11^ CFU per animal. Across the dose range tested, LS57 produced no significant effects on gastric acid secretion (including gastric juice pH), bile secretion, or gastrointestinal propulsion (Table 5).

**Table 5.** Safety pharmacology and oral toxicity studies of *E. ludwigii* LS57 in SD rats and Beagle dogs.

| **Study** | **Animal** | **Dose Levels (CFU/animal)** | **Key Findings** | **NOAEL (CFU/animal)** | **MTD (CFU/animal)** |
|---|---|---|---|---|---|
| Gastric acid secretion (Safety pharmacology) | SD rat | 1.2×10<sup>10</sup><br>1.2×10<sup>11</sup><br>6.0×10<sup>11</sup> | No significant effect on gastric acid secretion or pH | – | – |
| Bile secretion (Safety pharmacology) | SD rat | 1.2×10<sup>10</sup><br>1.2×10<sup>11</sup><br>6.0×10<sup>11</sup> | No significant effect on bile secretion | – | – |
| Gastrointestinal propulsion (Safety pharmacology) | SD rat | 1.2×10<sup>10</sup><br>1.2×10<sup>11</sup><br>6.0×10<sup>11</sup> | No significant effect on gastrointestinal propulsion | – | – |
| Single-dose oral toxicity | SD rat | 1.2×10<sup>10</sup><br>1.2×10<sup>11</sup><br>6.0×10<sup>11</sup> | Well tolerated; no mortality at the highest dose tested | – | 6.0×10<sup>11</sup> |
| Single-dose oral toxicity | Beagle dog | 1.2×10<sup>11</sup><br>4.7×10<sup>11</sup><br>2.1×10<sup>12</sup> | Well tolerated; no mortality at the highest dose tested | – | 2.1×10<sup>12</sup> |
| 1-week repeat-dose oral toxicity | SD rat | 1.2×10<sup>10</sup><br>1.2×10<sup>11</sup><br>6.0×10<sup>11</sup> | No mortality or treatment-related abnormalities in any examined parameter | 6.0×10<sup>11</sup> | – |
| 1-week repeat-dose oral toxicity | Beagle dog | 1.2×10<sup>11</sup><br>4.7×10<sup>11</sup><br>2.1×10<sup>12</sup> | No mortality or treatment-related abnormalities in any examined parameter | 2.1×10<sup>12</sup> | – |

Oral toxicity was assessed in two single-dose and two 1-week repeat-dose studies in SD rats and Beagle dogs. The Maximum Tolerated Dose (MTD) after a single oral administration was 6.0×10^11^ CFU per animal in rats and 2.1×10^12^ CFU per animal in dogs. After 7 consecutive days of repeated administration in both species, no mortality or moribundity occurred, and clinical observations, body weight, food consumption, body temperature, ECG, blood pressure, ophthalmologic examination, hematology, coagulation parameters, serum biochemistry, urinalysis, organ weights (absolute and relative), gross pathology, and histopathology revealed no treatment-related abnormalities. The No-Observed-Adverse-Effect Level (NOAEL) was 6.0×10^11^ CFU per animal in rats and 2.1×10^12^ CFU per animal in dogs (Table 5). Both repeat-dose studies were conducted in compliance with the U.S. Food and Drug Administration (FDA) GLP regulations.

## Methods

### Isolation of Swarming Bacteria from Fecal Samples

Fecal samples were stored at −20 °C and thawed on ice for 1 h. Fecal samples were weighed and resuspended at 100 mg/mL in sterile PBS (pH 7.4). A 3 μL aliquot of the suspension was spotted onto 0.5% LB agar plates (5 spots/plate, 3 plates/sample), yielding 15 technical replicates. Plates were incubated inverted at 37 °C with 40% humidity and observed at 16, 24, and 48 h. Samples lacking swarming motility were excluded. For swarming-positive samples, two isolates were picked from the swarm edge and streaked onto 1.5% LB agar plates for purification. Following overnight incubation, single colonies were inoculated into 3 mL LB broth and shaken at 200 rpm and 37 °C to obtain pure cultures of swarming strains.

### Gram Staining

Bacterial smears were prepared from mid-log phase cultures and heat-fixed by passing the slides briefly through a flame. The slides were sequentially stained with crystal violet (1 min) and iodine solution (1 min). Decolorization was performed with 95% ethanol for 20-30 s, with the process terminated immediately once the purple runoff ceased. Subsequently, the smears were counterstained with safranin O (30-60 s). Stained preparations were examined under an oil immersion microscope (100× objective), where bacteria retaining purple were classified as Gram-positive and those stained pink/red as Gram-negative.

### Growth Curve Determination

Frozen stocks were revived by overnight culture in flasks at 37 °C and 200 rpm. An equal volume of the resulting stationary-phase culture was subsequently inoculated into fresh medium. Bacterial growth was monitored by measuring optical density at 600 nm (OD₆₀₀) at regular intervals using a UV spectrophotometer, and the growth curve was plotted based on these measurements.

### Swarming Motility Assay and Time-lapse Imaging

Solid surface swarming motility was assessed on 0.5% LB agar plates. A 3 μL aliquot of bacterial culture was spotted onto the plate and incubated at 37 °C with 40% relative humidity. For static analysis, plates were monitored at various time points (3-16 h) to determine the onset and progression of swarming. Concurrently, dynamic swarming behavior was captured using a time-lapse imaging system: spotted plates were placed on an incubation stage, and image acquisition was performed via MVS camera software for 6-16 h. Recording was terminated once the swarm front reached the edge of the plate, and the resulting footage was processed to visualize continuous motility.

### Strain Identification by 16S rRNA Gene Sequencing

Genomic DNA was extracted from an overnight LB culture of LS57 with HiPure Bacterial DNA Kit. The near full-length 16S rRNA gene was amplified with the universal primers 27F (5′-AGAGTTTGATCCTGGCTCAG-3′) and 1492R (5′-GGTTACCTTGTTACGACTT-3′), and the products were checked by agarose gel electrophoresis, purified and sequenced in both directions. The consensus sequence was compared with the EzBioCloud 16S rRNA database^35^ and with the type strains of the *E. cloacae* complex.

### Whole-Genome Sequencing and Annotation

Genomic DNA was extracted from cell pellets using a CTAB-based protocol involving liquid nitrogen homogenization, phenol-chloroform-isoamyl alcohol (25:24:1) extraction, and isopropanol precipitation. The genome of strain LS57 was sequenced using a hybrid approach combining the DNBSEQ platform (BGISEQ-500, paired-end 150 bp; short reads) and the Oxford Nanopore PromethION platform (SQK-LSK114 library; long reads). Raw sequencing data were quality-controlled using Trimmomatic (filtering adapters, trimming low-quality bases <Q20, and removing reads <75 bp). Hybrid genome assembly was performed using Unicycler, integrating short-read data (SPAdes) with long-read data (miniasm/Racon) to resolve repetitive regions. Assembly quality was evaluated by mapping reads back to the consensus sequence to assess GC-content and coverage depth distribution. Coding genes were predicted using GeneMarkS, while non-coding RNAs were identified using RNAmmer and tRNAscan-SE. Prophage-like regions were identified with Prophage Hunter^55^ and confirmed with PHASTEST^56^, and insertion sequences were annotated with ISFinder. Functional annotation was performed by comparing predicted proteins against public databases, including GO^41^, KEGG^42^, CARD^28^, and VFDB^58^.

### Metabolite Profiling by LC-MS/MS

Metabolite extraction was performed using an automated workstation. Briefly, 50 μL aliquots of each sample (triplicate cultures of LS57 and LB controls) were combined with 200 μL ice-cold extraction solvent (methanol:acetonitrile, 1:1, v/v) containing internal standards. The mixtures were vortexed at 750 rpm for 5 min, left to stand for 5 min, and then filtered; the filtrate was collected for analysis. Polar metabolites were separated using a Vanquish UHPLC system coupled with an Orbitrap Exploris 120 mass spectrometer. Separation was performed on a Waters ACQUITY UPLC BEH Amide column (2.1 mm × 50 mm, 1.7 μm) maintained at 4 °C with an injection volume of 2 μL. The mobile phase consisted of (A) 25 mmol/L ammonium acetate and 25 mmol/L ammonia in water and (B) acetonitrile. MS data were acquired under Xcalibur control using electrospray ionization in positive (3.8 kV) and negative (−3.4 kV) modes. Key parameters included sheath gas 50 Arb, auxiliary gas 15 Arb, capillary temperature 320 °C, full MS resolution 60,000, MS/MS resolution 15,000, and stepped normalized collision energy (SNCE) values of 20, 30 and 40. Raw data were converted to mzXML using ProteoWizard (v3.0.24054)^59^ and imported into a custom R package for metabolite identification against an in-house database.

### DSS-Induced Colitis Model

Female C57BL/6J mice (8 weeks old) were provided with 2.5% DSS^45^ in drinking water ad libitum for 12 days to induce colitis. Mice were assigned to three groups (No DSS+LB, DSS+LB, and DSS+LS57) and were orally gavaged daily with either the test strain LS57 (4.0×10^9^ CFU/mL; 200 μL) or sterile LB medium (control). The experiment was terminated when the control group exhibited a body weight loss exceeding 20%, followed by euthanasia. Body weights were monitored daily throughout the study. Fecal samples were collected at baseline (day 0) and on day 12. Upon necropsy, colon lengths were measured, and tissues were harvested for multi-omics analysis. For histopathology, colons were flushed with PBS, infused with 4% PFA, rolled into “Swiss rolls,” fixed in 4% PFA at 4 °C overnight, and processed for H&E staining. For molecular analyses, colon tissues were snap-frozen in liquid nitrogen. Total RNA was extracted using Trizol for RT-qPCR analysis of pro-inflammatory cytokines (TNF-α, IL-1α, IL-1β, CXCL-1, and CCL-2) and the anti-inflammatory cytokine IL-10. Total protein was extracted using RIPA lysis buffer containing protease inhibitors, quantified via BCA assay, and used for downstream experiments such as ELISA.

### Fecal Microbiota Profiling by 16S rRNA Gene Sequencing

Genomic DNA was extracted from fecal samples using the HiPure Stool DNA Kit. The V3-V4 hypervariable regions of the bacterial 16S rRNA gene were amplified from 20-50 ng of genomic DNA as template, using the forward primer containing the sequence 5 ′-ACTCCTACGGGAGGCAGCAG-3 ′ and the reverse primer containing the sequence 5 ′-GGACTACHVGGGTWTCTAAT-3 ′. The first-round PCR was performed with an initial denaturation at 94 °C for 3 min, followed by 14-16 cycles of denaturation at 94 °C for 10 s, annealing at 53 °C for 30 s, and extension at 72 °C for 30 s, with a final extension at 72 °C for 5 min. Index adapters were then added to the amplicons via a second-round PCR using the first-round products as templates; the program included initial denaturation at 94 °C for 3 min, 10-12 cycles of denaturation at 94 °C for 10 s, annealing at 60 °C for 30 s, and extension at 72 °C for 15 s, with a final extension at 72 °C for 5 min. The final indexed libraries were purified, and fragment sizes (∼600 bp) were verified by electrophoresis on a 1.5% agarose gel. Library concentrations were quantified using a microplate reader (Tecan, Infinite 200 Pro). Finally, paired-end sequencing (PE250/PE300) was performed on an Illumina MiSeq/NovaSeq platform according to the manufacturer’s instructions to generate raw reads for 16S rRNA community profiling.

### TNBS-Induced Colitis Model

A 2% TNBS^49^ in 40% ethanol solution was prepared by thoroughly mixing 5% TNBS, absolute ethanol, and physiological saline at a ratio of 2:2:1. This solution was administered rectally to mice fasted for 16 h (day 0). Mice in the TNBS+LS57 group were gavaged daily for 4 consecutive days (days 0-3) with LS57 (7.0×10^8^ CFU per mouse), while TNBS+LB controls received an equal volume of LB medium. Body weight was monitored daily, and the experiment was terminated when the average body weight of the control group recovered to approximately 95% of baseline value. Mice were euthanized and colon length was measured.

### Co-administration of CsA and LS57 in DSS-Induced Colitis

A mouse colitis model was established using 3% DSS in drinking water (lasting 11 days). CsA (15 mg/kg) was administered daily by gavage to mice in the CsA group (DSS+CsA), while the combination therapy group (DSS+CsA+LS57) received both CsA (15 mg/kg) and LS57 bacterial suspension (8.0×10^8^ CFU/mouse) daily by gavage. Mice in the control group (DSS+Vehicle+LB) were gavaged with an equal volume of vehicle plus LB medium. Body weight was monitored daily, and the experiment was terminated when body weight in the control group decreased by 20%, after which colon length was measured.

### Construction of the Bioluminescent Strain LS57-LUC

SZU195 is a helper strain harboring plasmid pJMP1039 (Addgene #119239). This plasmid contains the *tnsABCD* operon and an ampicillin resistance marker, providing the necessary transposase machinery for Tn7-based transposition^60^. SZU215 is a tagging strain carrying a plasmid that comprises a gentamicin resistance gene, the left and right arms of the Tn7 transposon, and the *luxCDABE* reporter gene expression cassette driven by the strong promoter P3^61, 62^. LS57 (recipient), SZU195 (helper), and SZU215 (donor) were inoculated into 5 mL of LB and grown overnight (16 h) at 37 °C with 200 rpm shaking. LB for SZU195 was supplemented with 100 μ g/mL 2,6-diaminopimelic acid (DAP) and 100 μ g/mL ampicillin, and LB for SZU215 with 100 μ g/mL DAP and 20 μ g/mL gentamicin. Cells were harvested, washed twice with PBS, and resuspended in 100 μL LB. The three strains were mixed at a 1:1:1 ratio, spotted onto LB agar containing DAP, and incubated at 37 °C for 16 h for conjugation. The bacterial lawn was then scraped, resuspended in 1 mL LB, washed twice with PBS, and resuspended in 500 μL LB; 500 μL was plated onto LB agar containing 20 μg/mL gentamicin but lacking DAP to select transconjugants. Positive clones were verified by colony PCR and stored in 20% (v/v) glycerol at −80 °C.

### *In vivo* Bioluminescence Imaging and Biodistribution Analysis

A total of 25 C57BL/6J mice were enrolled and stratified by body weight into five groups (n=5 per group) to ensure no significant baseline weight differences. Mice were administered 200 μL of either LS57-LUC (∼4.0×10^9^ CFU/mL) or sterile LB broth via oral gavage. Bioluminescence imaging was performed using an AniView100 imaging system. Mice were anesthetized with isoflurane and imaged at specific time points post-gavage: Group 0 (LB control) was imaged immediately (0 h), while Groups 1-4 (LS57-LUC) were imaged at 3, 6, and 24 h. Following imaging, mice were euthanized, and gastrointestinal tissues (stomach, small intestine, cecum, and colon) were harvested, arranged on black card paper, and imaged *ex vivo* to assess bacterial biodistribution.

### Simulated Gastric Fluid Tolerance Assay

Acid and gastric fluid tolerance of stationary-phase LS57 cells was assessed. Overnight cultures were sub-cultured 1:100 into fresh LB medium and grown to log phase. Cells were harvested (4,000 × g, 5 min), resuspended in SGF adjusted to pH 2.0, 3.0 or 4.0, with cells held in 0.9% NaCl (pH 7.0) as the control^63^, and incubated at 37 °C with shaking (200 rpm). Aliquots were collected at hourly intervals from 0 to 4 h, serially diluted, and plated onto non-selective agar. After incubation, viable counts were enumerated and survival rates calculated relative to the initial inoculum.

### Cytotoxicity of LS57 in HT-29 Co-culture

The cytotoxic effects of LS57 on HT-29 were evaluated using a co-culture system. HT-29 cells were maintained in McCoy’s 5A medium supplemented with 10% fetal bovine serum and 1% penicillin/streptomycin. For the assay, cells were seeded into 96-well plates and cultured overnight to allow adhesion. Bacterial strains were revived in LB medium via overnight culture at 37 °C and 200 rpm. The following day, the saturated cultures were sub-cultured 1:100 into fresh LB medium and grown for an additional 3 h to reach a concentration of approximately 5.0×10^9^ CFU/mL. Serial dilutions of the bacterial suspension were then added to the adhered HT-29 cells to achieve final concentrations of 5.0×10^4^ and 5.0×10^5^ CFU/mL. Control groups included cells treated with sterile LB medium alone and untreated cells. Following a 24-h co-incubation period at 37 °C, cell viability was quantified using the Cell Counting Kit-8 assay according to the manufacturer’s instructions.

### Biofilm Formation Assay

Biofilm formation was quantitatively assessed using a 96-well peg lid system. Briefly, 180 μL of bacterial suspension was added to each well of a flat-bottomed 96-well plate, and the plate was sealed with a 96-peg lid. The assembly was incubated at 37 °C for 24 h to allow biofilm formation on the peg surfaces. For biofilm inhibition assays, bacterial suspensions were supplemented with the inhibitory agent (ε-poly-L-lysine, 0-2000 μg/mL) prior to inoculation. Following incubation, the peg lids were gently rinsed twice in sterile saline to remove non-adherent planktonic cells. Adherent biofilms were fixed with 0.1% Crystal Violet (CV) solution for 30 min at room temperature and subsequently washed twice more with saline. Bound CV was then solubilized using 95% ethanol, and subsequently, the absorbance of the eluted dye was measured at 590 nm using a microplate reader to quantify biofilm biomass. Biofilm formation was classified against a cut-off ODc defined as the mean A590 of the negative control plus three standard deviations, and strains with an A590 at or below ODc were recorded as non-biofilm-forming^51^.

### Hemolysis Assay

Hemolytic activity was evaluated on 5% defibrinated sheep blood agar plates. Sterile LB liquid medium was cooled to approximately 50 °C, supplemented with sterile defibrinated sheep blood to a final concentration of 5%, and poured into Petri dishes. Once solidified, bacterial suspensions were streaked onto the plates and incubated at 37 °C with 40% humidity for 24 h. Plates were subsequently observed for hemolysis, and representative images were documented.

### Disk Diffusion

Antimicrobial susceptibility was evaluated using the Kirby-Bauer disk diffusion^52^ method on Mueller-Hinton Agar (MHA). Overnight bacterial cultures grown in Tryptic Soy Broth (TSB) were adjusted to a 0.5 McFarland standard (equivalent to 1.5×10^8^ CFU/mL) and uniformly swabbed onto MHA plates. Antibiotic disks were applied aseptically, ensuring a minimum distance of 24 mm between disk centers and 15 mm from the plate edge. Plates were incubated inverted at 37 °C with 40% humidity for 16-18 h. Inhibition zones were measured using vernier calipers under transmitted light, interpreted according to the CLSI M100 standard^64^.

### Minimum Inhibitory Concentration (MIC)

Broth microdilution assays were performed in 96-well plates to determine MIC^53^ values. Two-fold serial dilutions of antibiotics were prepared in TSB. Each well was inoculated with an equal volume of standardized bacterial suspension (adjusted to a 0.5 McFarland standard) to achieve a final concentration of approximately 5.0×10^5^ CFU/mL. Plates included positive (bacteria only) and negative (medium only) controls. Following incubation at 37 °C for 18-24 h, bacterial growth was assessed by measuring the optical density at 600 nm (OD₆₀₀) using a microplate reader. The MIC was defined as the lowest antibiotic concentration that inhibited visible bacterial growth.

### *In Vitro* Conjugation Assay

*Escherichia coli* MG1655 transformed with pUC57-Tac-mCherry-Gen (gentamicin resistance) was used as the recipient strain, which grew on LB agar containing GEN. *E. coli* Stbl3 transformed with the RP4 plasmid (ampicillin/kanamycin/tetracycline resistance) was used as the Positive Control (PC) donor strain, which grew on LB agar containing AMP+KAN+TET. The donor (PC or LS57) and recipient strains were cultured in LB broth at 37 °C and 200 rpm for 16 h, diluted 1:100, and then cultured for an additional 3 h until the exponential growth phase. Subsequently, the mixed suspension of the PC donor and recipient (1:1) was inoculated onto LB agar containing AMP+KAN+TET+GEN or GEN alone, and cultured for 3 h at 37 °C and 40% humidity. Mixtures of LS57 and the recipient (1:1 or 1:10) were inoculated onto LB agar containing DOX+GEN or CMZ+GEN, and cultured for 3 h at 37 °C and 40% humidity. Antibiotic abbreviations and concentrations are as follows: GEN (gentamicin, 50 μg/mL), AMP (ampicillin, 100 μg/mL), KAN (kanamycin, 50 μg/mL), TET (tetracycline, 20 μg/mL), DOX (doxycycline, 10 μg/mL), CMZ (cefmetazole, 30 μg/mL).

### LS57 Translocation and Tissue Colonization Assay

To evaluate the potential of LS57 to cause bacteremia in both immunocompetent and immunodeficient hosts, C57BL/6J WT and IL-10 KO^21^ female mice (8-9 weeks old) were randomly assigned to four experimental groups: No DSS, DSS, DSS+EcN, and DSS+LS57. Mice in the DSS, DSS+EcN, and DSS+LS57 groups received 2.5% DSS in their drinking water together with daily oral gavage (600 μL), to induce colitis and disrupt the colonic barrier. After 10 consecutive days of oral administration of LS57 or EcN at 8.0×10^8^ CFU per mouse per day (or an equivalent volume of PBS for controls), tissues including colon, MLN, liver, and spleen were aseptically collected, weighed, and homogenized in an appropriate volume of PBS proportional to tissue weight. Each tissue homogenate was plated onto standard LB agar, LB agar containing 100 µg/mL cefazolin, and LB agar containing 200 µg/mL cefazolin to preliminarily detect the presence of LS57. In parallel, LB agar supplemented with 20 µg/mL spectinomycin was used to screen for EcN colonization. In addition, 100 µL of EDTA-K₂-anticoagulated whole blood was plated directly onto LB agar and incubated at 37 °C with 40% humidity for 16 h to assess bacterial translocation; cultures were extended to 24 h when no growth was evident at the initial reading. Positive control assays were performed by mixing 100 µL of EDTA-K₂- anticoagulated whole blood from healthy mice with 100 µL of diluted LS57 or EcN suspension (∼2.0×10^3^ CFU/mL) prior to plating under identical conditions. Colonies recovered from antibiotic-containing plates were isolated as pure cultures and identified by PCR to confirm whether they corresponded to LS57 or EcN.

### Antibiotic Treatment of LS57-Induced Bacteremia in Mice

Eight-week-old male and female C57BL/6J wild-type mice were randomly allocated into five groups per sex (*n*=7 per group) to evaluate the therapeutic efficacy of gentamicin (GM) and levofloxacin (Levo) against LS57–induced bacteremia. LS57 and EcN strains were cultured overnight in LB broth, sub-cultured to mid-log phase, washed twice with PBS, and resuspended in PBS to a final concentration of ∼2.0×10^8^ CFU/mL for intraperitoneal inoculation (500 μL per mouse). Disease progression was monitored at defined intervals (0, 1, 2, 20, 24, 44, 48, 68, 72, and 92 h post-inoculation) by measuring rectal temperature and applying MSS^57^. An MSS ≥ 3 or body temperature ≤ 35 °C was considered indicative of successful bacteremia induction, at which point antimicrobial treatment commenced. GM (62 mg/kg/24 h, intramuscular injection) or Levo (41 mg/kg/12 h, intravenous injection) was administered for up to 4 consecutive days, with 0.9% NaCl as the vehicle control. Blood, peritoneal lavage fluid, liver, spleen, and kidney samples were collected at 46 h and at the study endpoint (94 h), homogenized in PBS, and quantitatively cultured on LB agar plates to determine bacterial loads. Tissue histopathology was assessed using hematoxylin-eosin staining.

### Statistical Analysis

Comparisons between two groups used the two-tailed Mann-Whitney U test^65^. Comparisons among three or more groups (the DSS colitis, CsA combination and translocation experiments) used the Kruskal-Wallis test followed by Dunn’s multiple-comparison test^66^. In all figures, p < 0.05 is denoted as *, p < 0.01 as **, and p < 0.001 as ***. Animals were randomized by body weight before allocation, and histopathological scoring was performed by an observer blinded to group assignment, in accordance with the ARRIVE guidelines 2.0^67^.

ANI values were calculated with the OrthoANIu algorithm implemented in the EzBioCloud^35^ platform (https://www.ezbiocloud.net). 16S rRNA gene sequence similarity was queried against the same database, and species-level assignment followed the recommended 16S identity thresholds.

The dDDH values were calculated using the GGDC 3.0^38^ platform (https://ggdc.dsmz.de/home.php).

LEfSe^48^ analysis was performed with the LEfSe implementation of Segata et al. in Galaxy, using the factorial Kruskal-Wallis test (p < 0.05) and an LDA score threshold of 2.0 to identify differentially abundant taxa between groups.

Alpha diversity (Shannon index) was compared within each group between day 0 and day 12 by the Wilcoxon signed-rank test for paired samples, and between groups at a single time point by the Mann-Whitney U test. Metabolites were considered differentially abundant when the absolute log2 fold change exceeded 1.0 and the P value from Welch’s t test (Benjamini-Hochberg corrected) was below 0.05^68^.

### Bacterial Strains Used in This Study

The bacterial strains used in this study are listed in Table 6.

**Table 6.** Bacterial strains used in this study.

| Strain name | Description | Source |
|---|---|---|
| *Enterobacter ludwigii* LS57 | LBP candidate in this study | This study |
| *Enterobacter ludwigii* LS57-LUC | Bioluminescence imaging for *in vivo* biodistribution tracking in mice | This study |
| *Escherichia coli* ATCC 25922 | Positive control in biofilm formation assay | ATCC 25922<sup>69</sup> |
| *Staphylococcus aureus* LSU00019 | Positive control (α-hemolysis) in hemolysis assay | Lab collection |
| *Bacillus cereus* LS00095 | Positive control (β-hemolysis) in hemolysis assay | Lab collection |
| *Escherichia coli* MG1655 | Recipient strain in *in vitro* conjugation assay | ATCC 700926<sup>69</sup> |
| *Escherichia coli* Stbl3 | Positive control donor strain in *in vitro* conjugation assay | Study<sup>29</sup> |
| *Escherichia coli* Nissle 1917 (EcN) | Control strain in mouse translocation and bacteremia models | Study<sup>22</sup> |
| *Escherichia coli* SZU195 | Helper strain for LS57-LUC construction | Study<sup>60</sup> |
| *Escherichia coli* SZU215 | Donor strain for LS57-LUC construction | Study<sup>61, 62</sup> |

## Discussion

This study establishes a complete translational pipeline for the novel LBP *Enterobacter ludwigii* LS57, spanning from human-derived strain isolation to GLP-compliant preclinical evaluation. Most investigations of live biotherapeutics remain limited to *in vitro* assays or preclinical animal models, whereas the present work connects basic discovery to formulation development and GLP-compliant toxicology. Such a full-chain study is relatively rare in the microbiome field and provides a template for translating basic bacteriology into LBP development^18, 19^.

A key strength of this work lies in our function-first screening strategy. This work extends, and was inspired by, our previous work on *Enterobacter* sp. SM3, which is a mouse-isolated swarming strain that can ameliorate intestinal stress in mice^32^. We screened more than 100 fecal samples from human donors and selected LS57 first on the basis of its swarming phenotype, then confirmed its value by *in vivo* therapeutic efficacy and safety in mouse models, rather than on taxonomy alone. LS57 shares with SM3 its swarming motility, its protective effect in colitis and its membership of the genus *Enterobacter*, and therefore stands in the same relation to the human gut as SM3 does to the mouse gut. This approach ensures that LS57 is a human-gut-derived strain biologically suited to the human intestine and functionally relevant to intestinal disease, rather than a lab-adapted isolate with restricted therapeutic potential or a murine isolate that may only poorly colonize the human gut^70^.

LS57 is consistent with a multi-modal mechanism of action that distinguishes it from many conventional probiotics with unclear functional pathways. RT-qPCR revealed significant reductions in TNF-α, CXCL-1, and CCL-2 (Fig. 2D), and LS57 cultures were enriched for anti-inflammatory metabolites (Fig. 1I), supporting an immunomodulatory signature. Enrichment of *Faecalibaculum* and *Dubosiella* in LS57-treated mice (Fig. 2G) and temporal increases in SCFA-associated taxa align with improved mucosal homeostasis. Direct *in vivo* quantification of LS57-derived metabolites and mechanistic dependency on IL-10/Treg or AHR pathways were not tested here and remain to be established. The therapeutic effects of LS57 in multiple mouse models are associated with sustained intestinal colonization during repeated dosing, immunomodulatory activity of secreted metabolites, downregulation of pro-inflammatory cytokines, and a possible improvement of barrier integrity that was not directly measured here. Together, these features support stable and sustained biological effects in the gut microenvironment. However, one limitation of the colonization study is that the control bacterial strain used was *E. coli Nissle*, not an isogenic non-swarming mutant strain of LS57. Ideally, an isogenic mutant that abolishes swarming while preserving other phenotypes, such as swimming, growth, and surfactant secretion, would control for these variables. In future work, swarming-deficient mutants of LS57 will be generated to test the contribution of swarming motility to colonization and healing, and to define the mechanism involved.

Although *Proteobacteria* expansion is associated with IBD pathogenesis, this taxonomic association does not imply functional pathogenicity of all family members^71^. LS57 lacks the virulence determinants (*e.g.*, AIEC-defining adherence/invasion loci) characterizing disease-associated *Enterobacteriaceae*, exhibits γ-hemolysis, shows no systemic translocation in immunocompetent or immunodeficient hosts, and acts via a swarming-driven mechanism that presumably lowers luminal oxygen and expands beneficial anaerobes—effects that oppose, rather than recapitulate, the dysbiotic state. Consistent with this non-pathogenic, host-compatible profile, a separate mouse-derived *E. ludwigii* strain was independently shown to protect DSS-colitis via choline-mediated α7nAChR-RA/TGF-β signaling that promotes CD103^+^DC-induced Treg tolerance^43^, indicating that distinct *E. ludwigii* isolates can converge on anti-inflammatory outcomes through complementary ecological and immunometabolic routes.

A major highlight of the work is the comprehensive safety evaluation that directly addresses core potential regulatory concerns for LBPs. We performed antimicrobial susceptibility testing, MIC determination, genomic screening for virulence determinants, and bacterial translocation assessment in animal models. LS57 was sensitive to multiple antibiotics, exhibited low systemic translocation risk, and could be effectively eliminated if translocated into circulation, supporting a favorable safety profile. Translocation of bacteria into the colonic tissue and the MLN is an expected consequence of barrier disruption in colitis, and in our study both strains were recovered from these sites without reaching the blood, liver or spleen.

We further advanced LS57 from laboratory bacterial suspensions to a well-characterized drug product manufactured under cGMP conditions. GLP-compliant 1-week repeated-dose toxicity studies were conducted in rats and Beagle dogs, with comprehensive assessments of clinical signs, body weight, food consumption, body temperature, electrocardiography, blood pressure, ophthalmology, hematology, coagulation, serum biochemistry, urinalysis, organ weights, and histopathology. No significant adverse effects were detected, providing critical preclinical safety data to support human administration. A 90-day repeat-dose toxicity study in rats and dogs is ongoing to support a 3-month clinical trial for IBD patients.

Finally, the preclinical package presented here provides the basis for the first-in-human evaluation of LS57, which will be reported separately. With efficacy, safety, and manufacturing established, LS57 is positioned as a promising LBP candidate for intestinal inflammatory disorders.

Recent years have witnessed rapid development of LBPs for IBD and other microbiome-associated disorders. Several products, including Rebyota (RBX2660)^72^ and Vowst (SER-109)^73^, have now been approved for recurrent *Clostridioides difficile* infection, illustrating the potential of microbiome-based interventions to modulate host immunity and restore intestinal homeostasis^74^. However, most current LBPs rely on complex bacterial consortia or spore-forming anaerobes, which often present challenges in manufacturing standardization, quality control, and mechanistic characterization^23^. In contrast, LS57 represents a distinct therapeutic paradigm. Rather than being selected based on taxonomic composition or metabolite production alone, LS57 was identified through its swarming phenotype, a functional bacterial behavior previously shown to promote intestinal healing by reducing luminal oxygen tension and facilitating the expansion of beneficial anaerobic microbiota. As a single-strain product, LS57 also offers advantages in process development, batch consistency, and regulatory assessment. Importantly, the present study establishes a complete translational pathway from strain discovery and mechanistic validation to CMC development and GLP toxicology evaluation. To our knowledge, this is among the first reports describing the complete preclinical translation of a swarming-based bacterial therapeutic, thereby expanding the repertoire of functional phenotypes that can be exploited for next-generation microbiome medicines. Unlike most currently developed LBPs that are defined by their taxonomic identity or community composition, LS57 was selected based on a host-beneficial behavioral phenotype. This phenotype-oriented discovery strategy may provide a complementary framework for future microbiome drug development, in which microbial functions rather than taxonomy serve as the primary criteria for therapeutic strain selection.

## Conclusion

In this study, we isolated and identified LS57 from more than 100 human fecal samples through function-first screening. LS57 exhibited consistent therapeutic efficacy in multiple mouse models of colitis, which was associated with its immunomodulatory metabolites, sustained intestinal colonization, and reduced inflammatory responses. Comprehensive safety evaluations, including *in vitro* antimicrobial susceptibility testing, virulence gene analysis, *in vitro* conjugation, and *in vivo* translocation and systemic infection assessments, demonstrated that LS57 was sensitive to multiple antibiotics and showed no detectable conjugative transfer of resistance genes. Under cGMP conditions, we established a scalable manufacturing process and developed a stable, colon-targeted enteric-coated capsule filled with lyophilized LS57 powder, and subsequent GLP-compliant 1-week repeated-dose toxicity studies in rats and Beagle dogs revealed no treatment-related clinical, laboratory or histopathological findings, supporting its clinical safety profile. Collectively, these findings establish LS57 as a promising, safe, and mechanistically distinct LBP candidate, supporting its further clinical development for intestinal disorders.

## Declarations

### Ethics Approval

Human fecal samples were collected and used under protocols approved by the Ethics Committee of Ruijin Hospital, Shanghai Jiao Tong University School of Medicine (Approval No. 2023-292), the Medical Ethics Committee of Renji Hospital, Shanghai Jiao Tong University School of Medicine (Approval No. KY2021-168-B), and the Medical Ethics Committee of Zhongshan Hospital, Fudan University (Approval No. B2025-028R); written informed consent was obtained from all donors. All animal experiments were performed in accordance with the National Research Council’s Guide for the Care and Use of Laboratory Animals and are reported according to the ARRIVE guidelines. All animal procedures were approved by the Institutional Animal Care and Use Committee (IACUC) of the Laboratory Animal Center, Fudan University (Approval No. 2025-IMI-002). The safety assessment of LS57 in SD rats and Beagle dogs was approved by the IACUC of Beijing Joinn Biologics Co., Ltd. (Approval No. D25-S066-RD [S-ACU25-0449] for the rat study and Approval No. R25-S066-RD [S-ACU25-0450] for the dog study).

### Data Availability

The complete genome sequence of *Enterobacter ludwigii* LS57, its 16S rRNA gene sequence, and the raw sequencing reads have been deposited in GenBank and the Sequence Read Archive under BioProject PRJNA1559167 (BioSample SAMN64864137). All other data supporting the findings of this study are included in the published article and its supplementary information files.

### Author Contributions Statement

W.C. and L.L. conceived and designed the study and acquired funding. M.W. led the methodology, with contributions from W.C. and Z.Q. C.Z. curated the data, developed the software and prepared the figures. Y.G. performed the formal analysis, led the investigation and administered the project. W.C. and W.Y. validated the results. L.L. provided the main resources. Z.Q. contributed to conceptualization, investigation, methodology and supervision. Y.Z., S.G. and S.W. contributed to data curation and resources. L.F. contributed to conceptualization, investigation, validation and visualization. W.Y. contributed to formal analysis, investigation and resources. L.Z. critically revised the manuscript and prepared the revised submission files. W.C., C.Z., L.F. and L.L. wrote the main manuscript text. All authors reviewed and edited the manuscript.

## Acknowledgements

The authors thank Dr. Sridhar Mani for his consulting assistance during his tenure at Shanghai Lishan Biotechnology Co., Ltd. We are also grateful to Dr. Guangbo Ge, Dr. Zhijun Cao, Dr. Bing Su, and Dr. Jianpeng Ma for their insightful discussions, and to Dr. Xue Liu for preparing the fluorescent strains of LS57. In addition, we acknowledge Hangzhou Start Bioservices Biotechnology Co., Ltd. for CMC manufacturing and Beijing Joinn Biologics Co., Ltd. for toxicology studies. Lastly, we thank the Medical Science Data Center at Shanghai Medical College of Fudan University for providing technical support for our bioinformatic analysis.

