## Supplementary Material for "Preclinical Development of *Enterobacter ludwigii* LS57, a Human-Derived Swarming Strain, as a Live Biotherapeutic Product for Inflammatory Bowel Disease"

**Table S1.** Molecular mechanisms of antibiotic resistance genes in *E. ludwigii* LS57.

| Molecular mechanism of antibiotic resistance | Number of resistance genes |
|---|---|
| Antibiotic efflux | 101 |
| Target alteration | 64 |
| Reduced permeability to antibiotic | 17 |
| Loss of functional proteins | 8 |
| Antibiotic inactivation | 7 |
| Target protection | 4 |
| Target bypass | 2 |

**Note**. A single gene may be assigned to more than one mechanism; the 203 assignments in this table correspond to 180 unique genes.

**Table S2. Fragments of Insertion Sequences (ISs) detected in ARGs of *E. ludwigii* LS57.**

| IS name | IS identity% | Accession # | GeneID | Gene name | Gene identity% | ARO accession | Drug class |
|---|---|---|---|---|---|---|---|
| ISLhe9 | 48.8 | CP000517 | LBP57B.IS350000300 | ***Ecol_soxR_MULT*** | 92.763 | 3003381 | **cephalosporin;**fluoroquinolone; glycylcycline; penam; phenicol; rifamycin; **tetracycline** antibiotic |
| ISSsu9 | 35.6 | NC_012926 | LBP57B.IS350000712 | *dfrA3* | 53.125 | 3003105 | diaminopyrimidine antibiotic |
| ISCgl1 | 33.2 | U85507 | LBP57B.IS350000902 | *cmlv* | 36.269 | 3002700 | phenicol antibiotic |
| ISAba43 | 25.0 | NTFR0000000 | LBP57B.IS350001471 | ***Ecol_mdfA*** | 86.241 | 3001328 | **Tetracycline** antibiotic |
| IS5564 | 30.4 | AF024666 | LBP57B.IS350001970 | *cmlv* | 35.979 | 3002700 | phenicol antibiotic |
| ISBsp7 | 30.6 | FOML01000001 | LBP57B.IS350002235 | ***marA*** | 93.548 | 3000263 | **carbapenem;cephalosporin; cephamycin;** fluoroquinolone; glycylcycline;monobactam; penam; penem; phenicol; rifamycin; **tetracycline** antibiotic |

**Note.** The resistance or intermediate susceptibility of E. ludwigii LS57 to the drug classes shown in bold was verified by disk diffusion testing (Table 2).

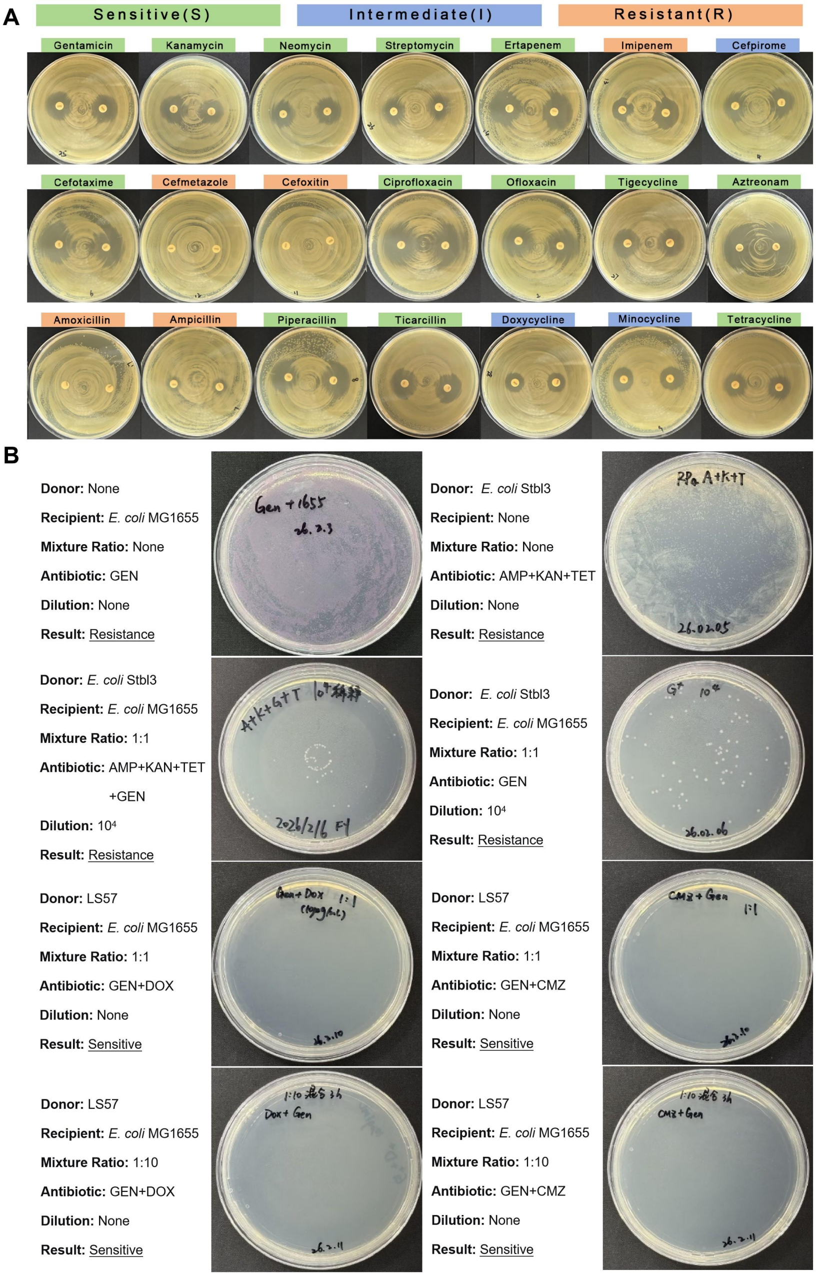


**Figure S1. *In vitro* antibiotic susceptibility and conjugative transfer potential of *E. ludwigii* LS57.** (A) Disk diffusion test on Mueller–Hinton agar, one antibiotic disk per plate; the coloured bar above each plate gives the interpreted category (green, susceptible; blue, intermediate; orange, resistant) according to CLSI M100 breakpoints. Zone diameters and the cognate resistance determinants are given in Table 2 and Fig. 4E. (B) Conjugation assay. *E. coli* MG1655 (pUC57-Tac-mCherry-Gen) was the recipient and LS57 the test donor, mixed at donor:recipient ratios of 1:1 and 1:10 and selected on gentamicin plus doxycycline (GEN+DOX) or gentamicin plus cefmetazole (GEN+CMZ). An RP4-carrying *E. coli* Stbl3 served as the positive-control donor (AMP+KAN+TET+GEN, dilution $10^{-4}$); the recipient and the donor plated alone on their respective antibiotics were growth controls. Transconjugants arose only in the positive control (65.1%, 175/269 recipient CFU); no colony was recovered from any LS57 mating (0/269), below the detection limit of the assay. Each panel lists donor, recipient, ratio, selection antibiotic and dilution together with the result (Resistance, growth; Sensitive, no growth). Antibiotic concentrations: GEN 50, AMP 100, KAN 50, TET 20, DOX 10 and CMZ 30 μg/mL.

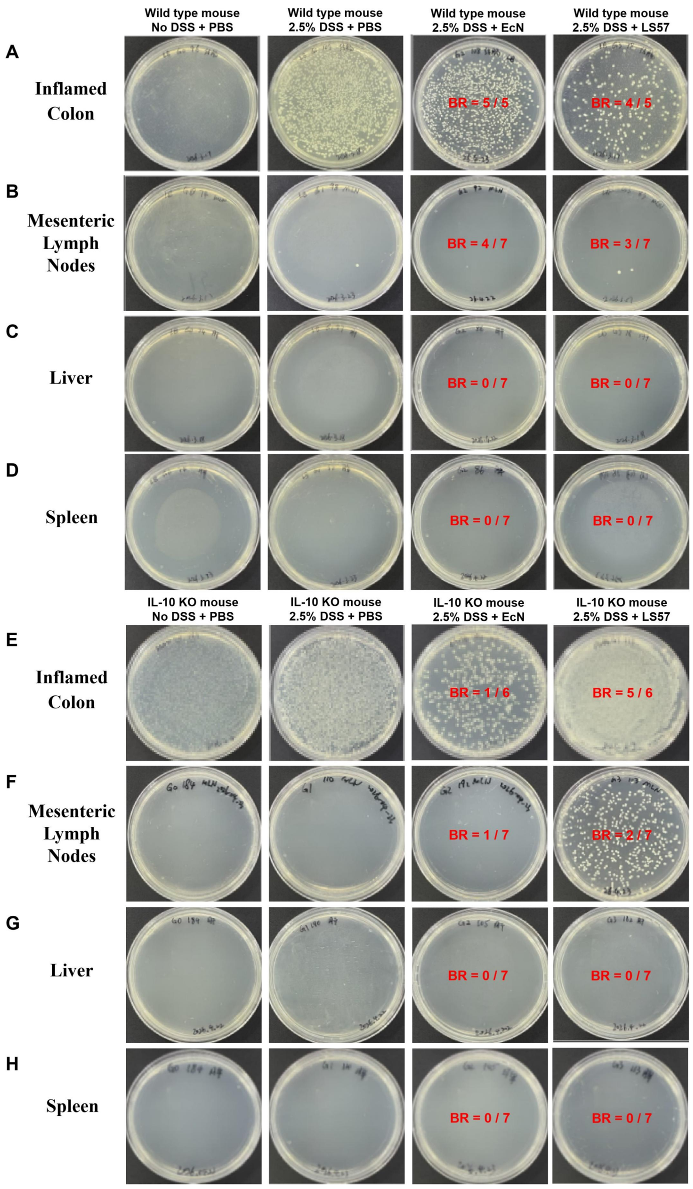


**Figure S2. Systemic translocation of *E. ludwigii* LS57 and *E. coli* Nissle 1917 (EcN) in DSS-induced colitis.** C57BL/6 wild-type (WT) and IL-10 knockout (IL-10 KO) mice received 2.5% DSS for 10 days and were gavaged daily with $8.0 \times 10^8$ CFU of LS57 or EcN, as in Fig. 5A, B; mice given no DSS plus PBS and mice given DSS alone served as controls. Colon, mesenteric lymph nodes, liver and spleen were cultured at necropsy, and one representative plate is shown for each condition (breakthrough ratios, culture-positive samples/total samples). (A) Representative cultures of the inflamed colon from WT mice; LS57 and EcN colonized the inflamed colon in 4/5 (80.0%) and 5/5 (100.0%) animals. (B) Representative cultures of mesenteric lymph nodes from WT mice; LS57 and EcN were recovered from 3/7 (42.9%) and 4/7 (57.1%) mice. (C) Representative cultures of liver from WT mice; neither strain was recovered (0/7). (D) Representative cultures of spleen from WT mice; neither strain was recovered (0/7). (E) Representative cultures of the inflamed colon from IL-10 KO mice; colonization was 5/6 (83.3%) for LS57 and 1/6 (16.7%) for EcN. (F) Representative cultures of mesenteric lymph nodes from IL-10 KO mice; LS57 and EcN were recovered from 2/7 (28.6%) and 1/7 (14.3%) mice. (G) Representative cultures of liver from IL-10 KO mice; neither strain was recovered (0/7). (H) Representative cultures of spleen from IL-10 KO mice; neither strain was recovered (0/7).

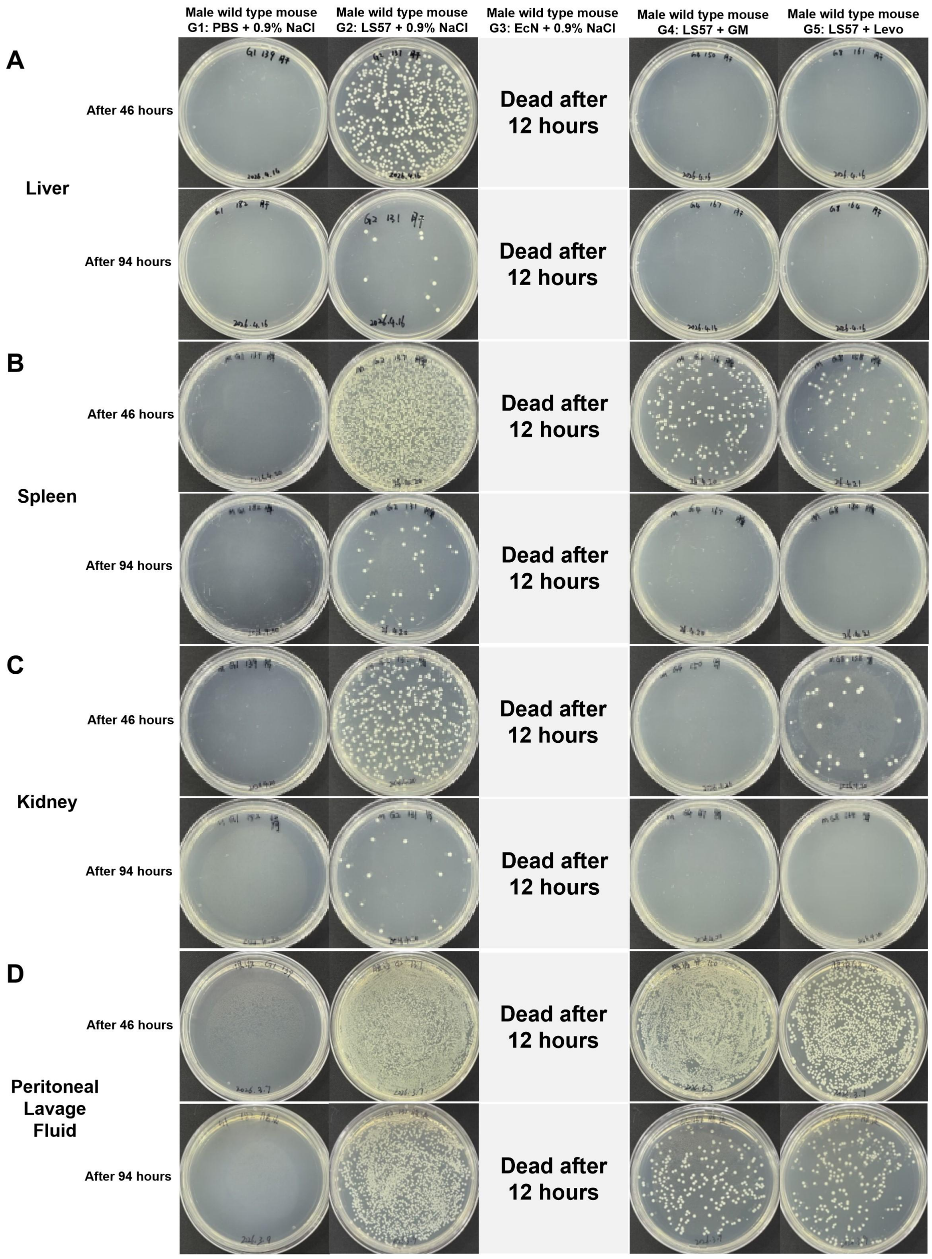


**Figure S3. Tissue and peritoneal clearance of *E. ludwigii* LS57 in male mice after antibiotic treatment.** Male wild-type (WT) mice were inoculated intraperitoneally with $1.0\times10^{8}$ CFU of LS57 and treated with gentamicin (GM) or levofloxacin (Levo) as in Fig. 5C, D (G1, PBS + 0.9% NaCl; G2, LS57 + 0.9% NaCl; G3, EcN + 0.9% NaCl; G4, LS57 + GM; G5, LS57 + Levo). Liver, spleen, kidney and peritoneal lavage fluid were sampled at 46 h (upper row of each pair) and 94 h (lower row) after infection; two plates are shown for each group and time point, except G3, in which all mice died within 12 h of EcN inoculation and a single plate is shown for both time points. (A) Representative cultures of liver; in antibiotic-treated animals (G4 and G5), colonies were still recovered in at least one group at 46 h, and the organ was culture-negative by 94 h. (B) Representative cultures of spleen; in antibiotic-treated animals, colonies were still recovered in at least one group at 46 h, and the organ was culture-negative by 94 h. (C) Representative cultures of kidney; in antibiotic-treated animals, colonies were still recovered in at least one group at 46 h, and the organ was culture-negative by 94 h. (D) Representative cultures of peritoneal lavage fluid; the lavage fluid of antibiotic-treated animals remained culture-positive at 94 h.

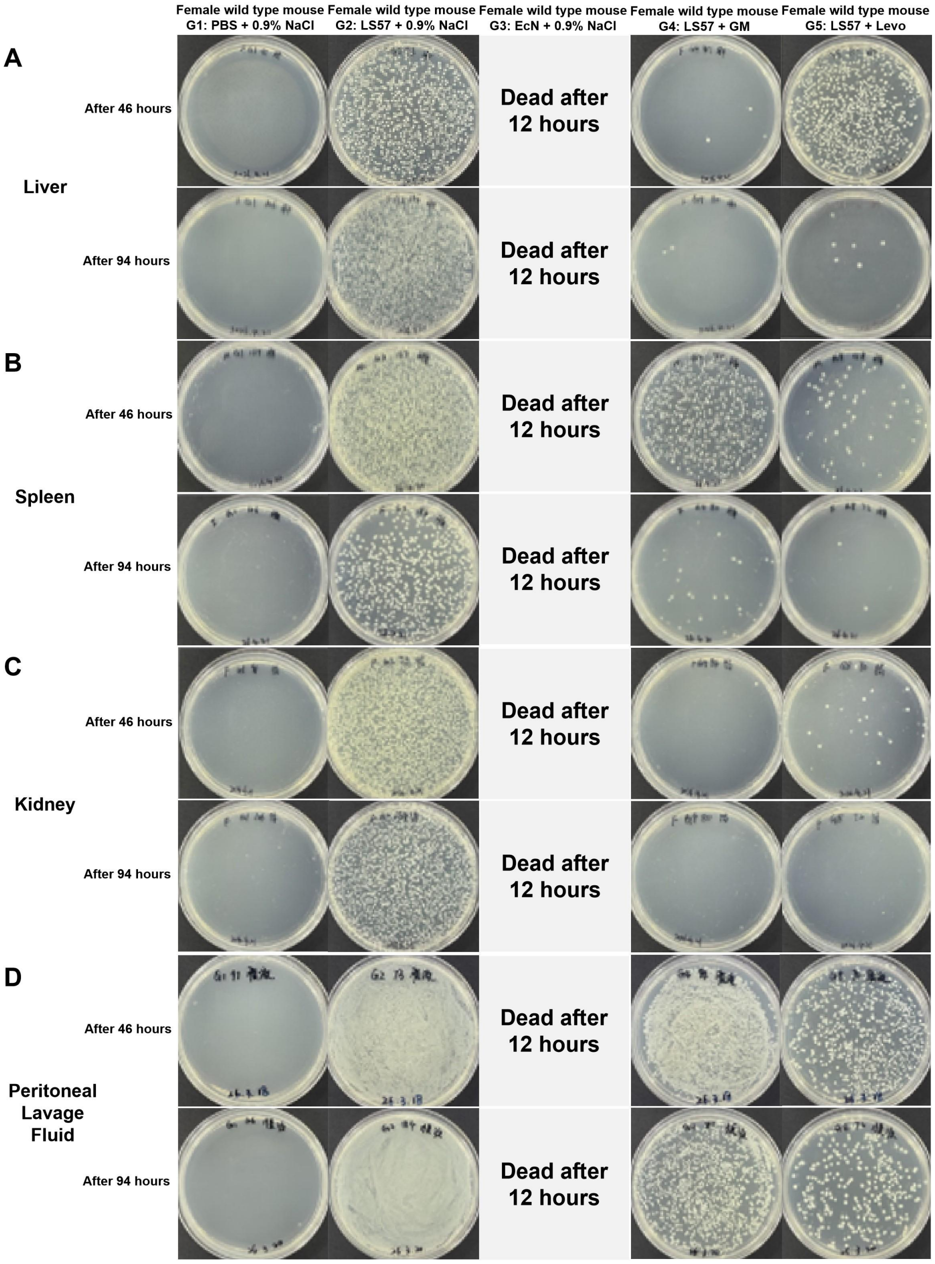


**Figure S4. Tissue and peritoneal clearance of *E. ludwigii* LS57 in female mice after antibiotic treatment.** Female WT mice were inoculated and treated as in Fig. S3 (G1, PBS + 0.9% NaCl; G2, LS57 + 0.9% NaCl; G3, EcN + 0.9% NaCl; G4, LS57 + GM; G5, LS57 + Levo). Liver, spleen, kidney and peritoneal lavage fluid were sampled at 46 h (upper row of each pair) and 94 h (lower row) after infection; two plates are shown for each group and time point, except G3, in which all mice died within 12 h and a single plate is shown for both time points. (A) Representative cultures of liver; in antibiotic-treated females (G4 and G5), colonies were still recovered in at least one group at 46 h, and low numbers of colonies persisted in both treatment groups at 94 h. (B) Representative cultures of spleen; in antibiotic-treated females, colonies were still recovered in at least one group at 46 h, and low numbers of colonies persisted in both treatment groups at 94 h. (C) Representative cultures of kidney; in antibiotic-treated females, colonies were still recovered in at least one group at 46 h, and the organ was culture-negative by 94 h. (D) Representative cultures of peritoneal lavage fluid; the lavage fluid of antibiotic-treated females remained culture-positive at 94 h.
